# Surface crosslinking of virus-like particles increases resistance to proteases, low pH, and mechanical stress for mucosal applications

**DOI:** 10.1101/2023.07.29.550271

**Authors:** Ahmed Ali, Suwannee Ganguillet, Yagmur Turgay, Tim Keys, Erika Causa, Ricardo Fradique, Viviane Lutz-Bueno, Serge Chesnov, Chia-wei Lin, Verena Lentsch, Jurij Kotar, Pietro Cicuta, Raffaele Mezzenga, Emma Slack, Milad Radiom

**Affiliations:** Institute of Food, Nutrition and Health, ETH Zürich, Zürich, Switzerland; Biological and Soft Systems, Cavendish Laboratory, University of Cambridge, Cambridge CB3 0HE, U.K; Paul Scherrer Institute PSI, Villigen, Switzerland; Laboratoire Léon Brillouin, CEA-CNRS (UMR-12), CEA Saclay, Université Paris-Saclay, 91191, Gif-sur-Yvette Cedex, France; University of Zürich/ETH Zürich, Functional Genomics Centre Zürich, Zürich, Switzerland

**Keywords:** Virus-like particle, Polyethylene glycol, Surface-crosslinking, Force spectroscopy, In vitro human nasal epithelial tissue, Mucus

## Abstract

Virus-like particles (VLPs) are emerging as nano-scaffolds in a variety of biomedical applications including the delivery of vaccine antigens to mucosal surfaces. These soft, colloidal, and proteinaceous structures (capsids) are nevertheless susceptible to mucosal environmental factors which limit their usefulness. We addressed this issue by crosslinking multiple capsid surface reactive residues using polyethylene glycol tethers. Surface crosslinking enhanced the colloidal stability and mechanical strength of VLPs against low pH, proteases, and mechanical agitation, while it did not interfere with function as vaccine. Chemical crosslinking thus offers a viable means to enhance the resilience of VLPs in mucosal applications.

## Introduction

Virus-like particles (VLPs) have emerged as invaluable nano-carriers in biomedical applications (*1,2*). Sensitive cargo can be protected by packaging it inside their capsid while genetic and/or chemical modifications can be used to deliver vaccine antigens or to target the cargo to specific cell types. In recent years, various therapeutic molecules have been integrated into VLPs for immunotherapy (*3*), gene therapy (*4–6*), chemotherapy (*7–9*), as well as for contrast imaging and photothermal therapy (*10,11*).

As carriers, VLPs must endure environmental challenges and it is likely that their material properties are an important factor determining their fate (*12*). General key considerations for “delivery-capable” VLPs include size, colloidal stability, target specificity, and responsiveness to stimuli for cargo release. For mucosal applications, VLPs are faced with additional physiological challenges including a thick and adhesive mucus gel, mucus regeneration and clearance, high concentrations of proteases and secretory immunoglobulin A (sIgA) antibodies and intermittent mechanical agitation (*13–15*). Together these effects can significantly reduce the efficacy of VLPs for mucosal applications (**Figure 1(a)**).

**Figure 1:**
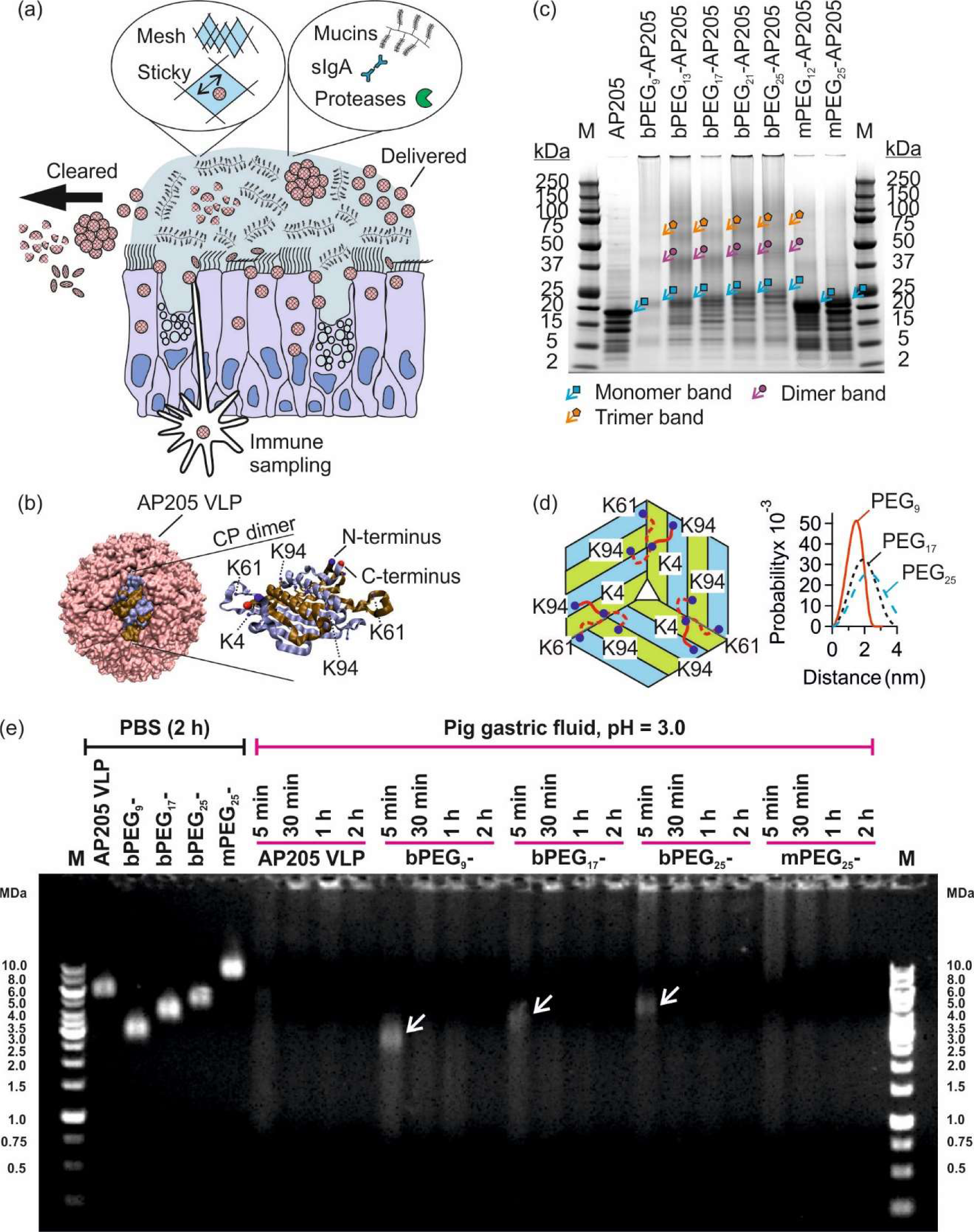
Susceptibility of virus-like particles (VLPs) to mucosal factors, and their capsid stabilization by means of surface crosslinking using macromolecule tethers. (a) VLPs are exposed to physical and chemical challenges on mucosal surfaces. (b) Schematic of AP205 VLP and schematic of one coat protein (CP) dimer (*41*). The N-terminus (blue) of one monomer, the C-terminus (red) of the other monomer, together with several lysine residues are marked. (c) Reducing SDS polyacrylamide gel electrophoresis of native, PEG-crosslinked (bPEG_n_-, n = 9–25), and PEGylated but not crosslinked AP205 VLPs (mPEG_n_-, n = 12 or 25). M is PageRuler Plus Prestained Protein Ladder. The monomer region is marked with cyan arrows/squares, dimer region with purple arrows/circles and trimer region with orange arrows/pentagons. (d) Schematic of AP205 CP arrangement near three-fold axis of symmetry and the assignment of lysine residues K4, K61 and K94 that participate in crosslinking reaction. In the schematic, 3 dimers are shown with their monomers colored in blue and green. Only one tether (red coil), that is either of the two of K4–K94 possibilities or the one of K4–K61 possibility, can occur. The end-to-end probability distributions of bPEG_n_ tethers (n = 9, 17 and 25) calculated from an analytical worm-like chain model (*43*). (e) Agarose gel electrophoresis of AP205 VLP, bPEG_9_-AP205 VLP, bPEG_17_-AP205 VLP, bPEG_25_-AP205 VLP and mPEG_25_-AP205 VLP in PBS and in pig gastric fluid (pH 3.0), respectively on the left and right of the gel. Incubation times varied between 5 min, 30 min, 1 hour and 2 hours at 37°C. Arrows at time point 5 min indicate persistent PEG-crosslinked AP205 VLPs in gastric fluid. M stands for marker.

To this end, there is compelling evidence that viruses, viral vectors, and VLPs disintegrate under mechanical force (*16–25*), and in unfavorable chemical environments (*26,27*). This raised the question: can surface crosslinking using macromolecule tethers enhance the stability of VLP capsids? To address this question, we employed homobifunctional polyethylene glycol (PEG) to tether surface lysine residues on capsid surface. VLP or viral vector PEGylation, without crosslinking, has been exploited as a means to enhance biological function, especially with adenovirus vectors (Adv). For example, PEGylation protected Adv from neutralizing antibodies, prolonged transgene expression and reduced immune activation against the vector (*28–30*). Subsequent experiments revealed a complex dependence on bio-physicochemical properties (*31–33*). Thereby, for example, conjugation optimization led to reduced anti-vector serum antibody and more persistent transgene expression (*34*). Other macromolecules were conjugated to bacteriophage Q*β* VLP to shield it from immune recognition (*35,36*). In other directions, PEG was used to tether a fluorescent molecule and folic acid to cowpea mosaic virus for intravital vascular imaging and targeted cargo delivery to cancer cell lines, respectively (*37,38*).

Our model VLP is assembled from 180 copies of *Acinetobacter* phage coat protein AP205 (*25*). AP205 VLP is a versatile platform for genetic and/or chemical conjugation of various antigenic molecules thus potentiating a vaccination strategy (*25, 39-41*). We investigated surface crosslinking of AP205 VLP using PEG tethers. To explore different modes of PEGylation, the AP205 VLP was reacted with homobifunctional (leading to surface crosslinking) and monofunctional (leading to simple PEGylation) linkers. We found that surface crosslinking is a viable toolbox that enhances the various stability profiles of VLPs, of which resilience to mechanical agitation, pH and protease stress are related to mucosal applications. Particularly, PEG crosslinking did not shield immune recognition of capsid coat proteins, enhancing vaccine stability without inhibiting immunogenicity.

## Results

Surface crosslinked and simply PEGylated VLPs were produced using homobifunctional and monofunctional PEG molecules respectively. The resulting VLPs are named bPEG_n_- and mPEG_n_-AP205 VLP where n is the number of PEG monomers. The functional group(s) of the PEG molecules was N-hydroxysuccinimide (NHS) ester which reacts with lysine residues and N-terminus of coat proteins (c.f. **Figure 1(b)**). The sequence of AP205 monomer and the details of the PEG molecules are presented in **S1**. We note that in the conditions used, the crosslinking was at single VLP level, and we did not observe aggregation between the particles (**S2**). Transmission electron microscopy (TEM) showed that the VLPs retained their spherical geometry after the reaction with the linker molecules and no detectable VLP size change for any lengths of PEG crosslinker tested, n = 9–25, was observed (**S3**).

### Crosslinking AP205 VLP capsid using polyethylene glycol (PEG) molecules

Investigations with mass spectrometry and reducing SDS-polyacrylamide gel electrophoresis (SDS PAGE) revealed that AP205 VLP reaction with homobifunctional PEG crosslinkers resulted in PEG-crosslinked AP205 dimers, trimers, and possibly higher oligomers. In the representative SDS PAGE shown in **Figure 1(c)**, in the lanes associated with bPEG_n_-AP205 VLPs (n = 9 to 25), bands in the MW range 37–50 kDa (marked with purple arrows/circles) correspond to PEG-crosslinked AP205 dimers, and bands in MW range 50–75 kDa (marked with orange arrows/pentagons) correspond to PEG-crosslinked AP205 trimers. Using nano ultra-performance liquid chromatography coupled to mass spectroscopy, it was found that lysine (K) residues K4 and K61 and K4 and K94 were involved in the crosslinking reaction (**Table S4-7**). In **Figure 1(d)**, a schematic of PEG linkages between individual AP205 monomers near the three-fold axis of symmetry is shown. Using coordinates from the cryo-electron microscopy reconstruction of the AP205 VLP (5LQP) in Visual Molecular Dynamics (VMD) (*41,42*), the N-N interatomic distances in K4–K61 linkage were estimated to be 1.8 nm and in K4–K94 linkage to be 1.6 nm and 2.4 nm. Calculation of end-to-end probability distributions of PEG_9_, PEG_17_ and PEG_25_ linker molecules using worm-like chain model are shown in **Figure 1(d)** (*43*). The calculated extension lengths are consistent with the interatomic distances. Additional details regarding the conjugation stoichiometry, mass spectroscopy and SDS PAGE are found in **S2**, **S4** and **S5**, respectively.

### Enhanced colloidal and enzymatic stability of surface-crosslinked AP205 VLPs

VLPs are potentially interesting scaffolds for the oral or intranasal delivery of biomolecules, requiring stability in pH-varying environments and in the presence of high concentrations of proteases (*1, 2, 12*). Therefore, increasing their stability in the complex mucosal environment is of high translational relevance.

Experiments using dynamic light scattering (DLS) showed that native VLP was stable at pH 7.4 but aggregated at pH 2.0–6.0 (**Fig. S6-1**). PEG-crosslinking increased the stability of VLP down to pH ∼ 4.0 (**Fig. S6-2**). We then tested the enzymatic stability of PEG-crosslinked VLPs in pig and mouse gastric fluids. In PBS, an incubation time of 2 h (or longer) at 37°C did not affect the stability or electrophoretic mobility of the VLPs (*25*) (**Figure 1(e)**). In pig gastric fluid, which had a pH of 3.0 and high protease concentration, PEG-crosslinked VLPs, namely bPEG_9_-AP205 VLP, bPEG_17_-AP205 VLP, and bPEG_25_-AP205 VLP, showed improved stability after 5 min of incubation, i.e., a band, marked with arrow, remained at the expected MW, with fewer breakdown products than the unmodified VLP. At the later time points of 30 min, 1 h and 2 h, the intact portion was disintegrated, and signs of aggregation and accumulation of material in the wells of the gel appeared. Interestingly, the PEGylated but not crosslinked VLP, namely mPEG_25_-AP205 VLP, showed a similar behavior to native VLP and disintegrated in as little as 5 min. This observation indicates that PEG-crosslinking VLP capsid, but not PEGylation per se, is a viable approach to overcome the susceptibility of VLPs to colloidal and enzymatic instability.

Experiments in pig gastric fluid at pH 4.7 corroborated the stabilizing effect of PEG-crosslinking, but not simple PEGylation, in the presence of proteases (**Fig. S7-1**). At this pH, the PEG-crosslinked VLPs were stable at all time points. In pig gastric fluid at pH 5.5, all VLPs were found to be stable over 1 h of incubation (**Fig. S7-2**). Incubation in mouse gastric fluid at pH 4.6 presented a similar behavior where the PEG-crosslinked VLPs appeared to be more stable compared to the naked and PEGylated but not crosslinked VLP (**Fig. S7-3**).

### Enhanced stiffness of surface-crosslinked AP205 VLPs

Susceptibility to mechanical stress sets a limit on potential applications of VLPs in various indications. This occurs during interactions with cells, e.g. during endocytosis or antigen presentation to B cells, or when exposed to mucus or fecal flow (*44–50*). Therefore, we examined the nanomechanical properties of native and PEG-crosslinked VLPs using atomic force microscopy (AFM) force-indentation spectroscopy.

The schematic of AFM measurements is depicted in **Figure 2(a)** and details are provided in the methods section. The example shown in **Figure 2(b)** depicts a single representative AP205 VLP for which the height prior to indentation (*H*_i_) and after indentation (*H*_f_) were evaluated. **Figure 2(c)** shows the heights distributions of AP205 VLP with the respective average values. The force versus indentation distance (*F* − *δ*) curve was used to calculate the stiffness, and the force versus indentation time (*F* − *t*) curve to calculate the strain rate. An example of these curves for the same VLP (**Figure 2(b)**) is shown in **Figure 2(d)**. The linear part of the response was used to calculate the stiffness, *k* = *dF/dδ*, the total indentation, *δ* = Δ*F/k*, and the indentation force rate *Ḟ* = *dF/dt*, which was converted to strain rate via *γ* = *Ḟ/kδ*. After an initial linear response, the VLP yielded at a threshold force/indentation. The histograms of maximal force and indentation (indent.) of AP205 VLP together with the average values of these parameters are shown in **Figure 2(e)**. The physical significance of these parameters is that at a force (or indentation) higher than the average maximal force (or indentation), yield occurs which significantly reduces the original stiffness of the VLP (*16-18, 22-25*). The histograms of stiffness and strain rate of AP205 VLP together with the average values of these parameters are shown in **Figure 2(f) and (g)**, respectively. For this VLP, we previously showed that the standard linear solid model (SLSM, **Figure 2(h)**) interprets the stiffness as a function of the strain rate (*25*). We found that at a strain rate < 100 1/s, the overall mechanical response is governed by *k*_0_ in SLSM. We retrieved a good agreement between the current evaluation of AP205 stiffness (47±15 pN/nm) with the previous one (53±23 pN/nm) at low strain rates.

**Figure 2:**
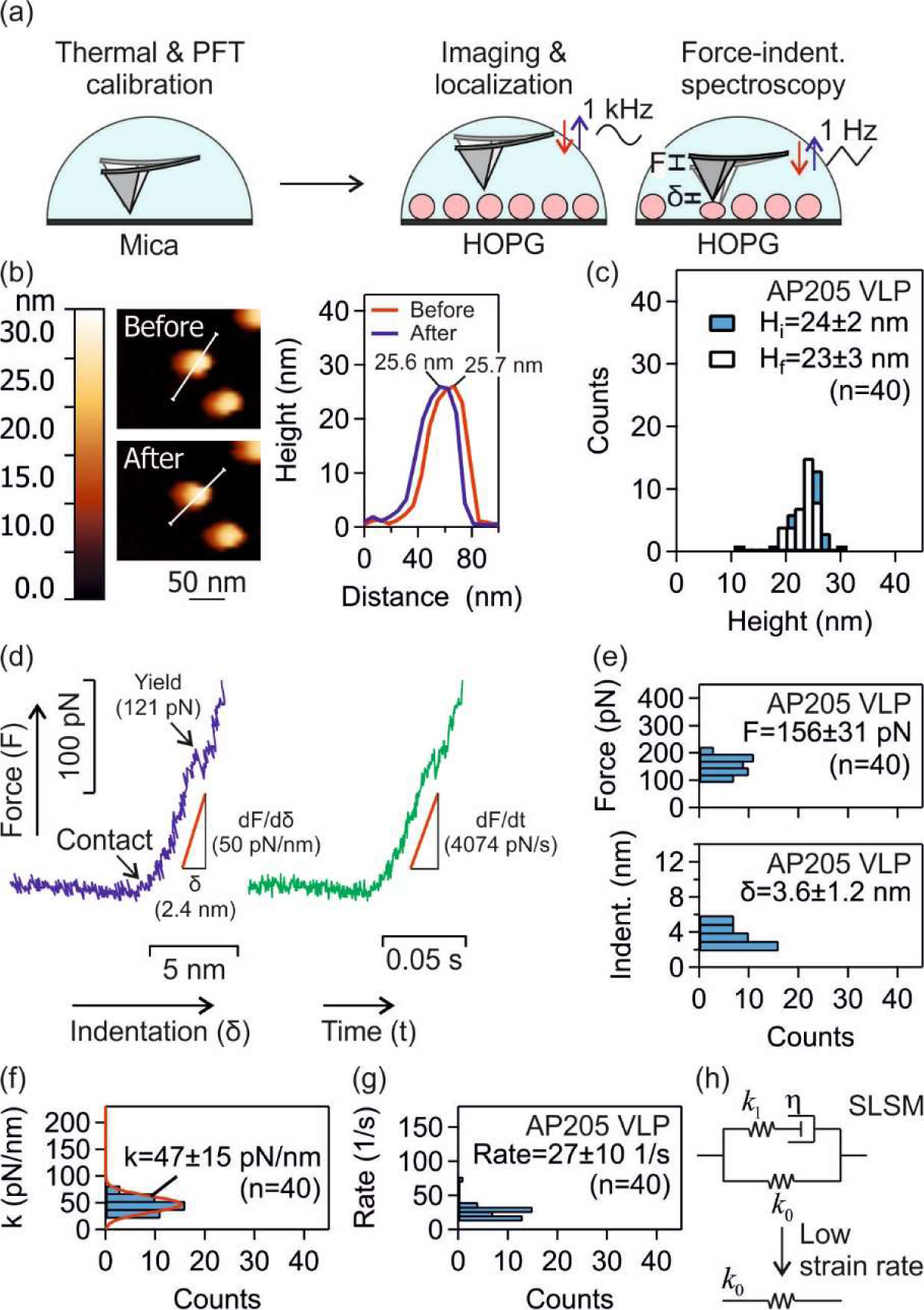
Mechanical properties of AP205 VLP investigated using AFM force-indentation spectroscopy. (a) Schematic of AFM calibration on freshly cleaved mica, and force-indentation spectroscopy of HOPG-immobilized VLPs. Single VLPs were localized using PeakForce tapping imaging (base tapping rate of 1 kHz) prior to force measurements using normal ramp at base approach-retraction rate at 1 Hz. (b) Example of height evaluation for a single AP205 VLP before and after force application. (c) Height distributions of AP205 VLP before and after force measurements. (d) Example of force-indentation distance and force-indentation time curves for AP205 VLP. The linear regime of force-indentation distance response was used to calculate the stiffness. In the same range, a linear fit to force-indentation time response gave the indentation force rate. (e, f and g) Histograms of maximal force and indentation (indent.) (e), stiffness (f) and strain rate (g) of AP205 VLP. Mean ± standard deviation is reported. The number (n) of AP205 VLPs in repeat measurements is indicated. (h) Standard linear solid model (SLSM). At low strain rates, there is no contribution from friction (or rate dependent resistance) to indentation.

Similarly, the mechanical properties of PEG-crosslinked VLPs, namely bPEG_9_-AP205 VLP, bPEG_17_-AP205 VLP, bPEG_25_-AP205 VLP were evaluated, and the results are shown in **Figure 3**. The initial heights and the heights after a single indentation showed similarity among the native and PEG-crosslinked VLPs. The distributions of maximal force and indentation (indent.) however showed variations compared to native VLP. The average maximal force was found to be higher and accordingly the average maximal indentation lower compared to native VLP. The increased (decreased) maximal force (indentation) of PEG-crosslinked VLPs indicated reduced susceptibility to mechanical agitation. Accordingly, the average stiffness for these VLPs was also higher by a factor of 1.3 to 2.0 than the stiffness of native VLP at similar range of strain rate.

**Figure 3:**
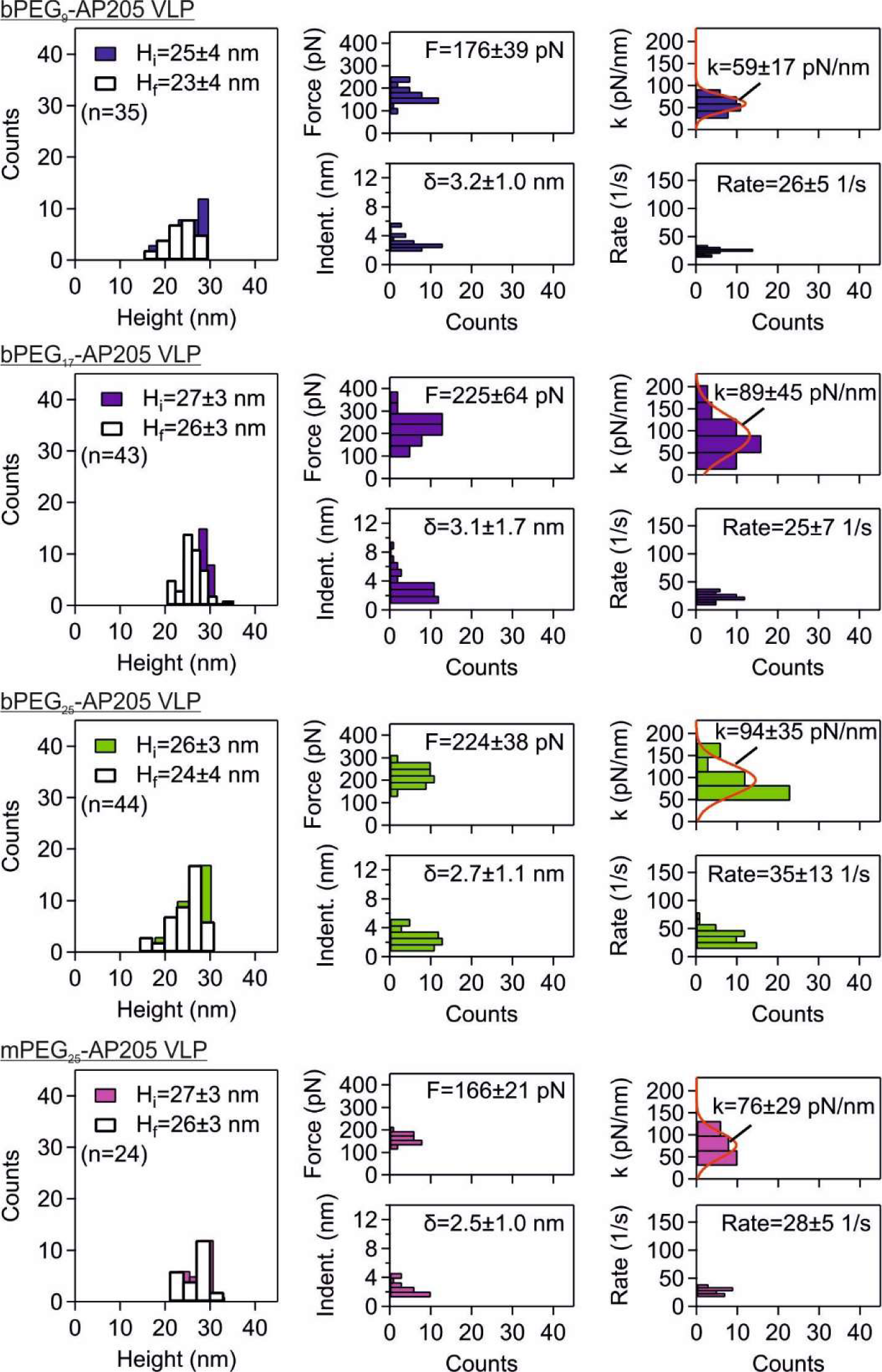
Enhanced strength and stiffness of PEG-crosslinked AP205 VLPs. Mechanical properties of PEG-crosslinked AP205 VLPs, namely bPEG_9_-AP205 VLP, bPEG_17_-AP205 VLP and bPEG_25_-AP205 VLP, and PEGylated but not crosslinked mPEG_25_-AP205 VLP investigated using AFM force-indentation spectroscopy. Distributions of height before and after indentation, maximal force/indentation in linear response regime, and stiffness/strain rate are shown together with mean ± standard deviation. The number (n) of VLPs in repeat measurements is indicated.

We then evaluated capsid size and shell thickness using small angle X-ray scattering (SAXS). The X-ray scattering spectra of the native and PEGylated VLPs together with fits to spherical core-shell form factor are shown in **Figure 4(a)**. In each case, we found a good agreement based on the low values for the fitting error (*χ* ^2^). In the summary panel in **Figure 4(b)**, it is shown that the diameter (*D*) and shell thickness (*h*) of VLP increased after PEGylation. In addition, while the VLP diameter remained somewhat constant between the different lengths of PEG tethers, the shell thickness increased with PEG length and had its highest value for the PEGylated but not crosslinked VLP. Subsequent to these evaluations, the elastic modulus was calculated from the relation *E* = *Dk*/2*h*^2^, **Figure 4(b)** (*51,52*). The elastic moduli remained somewhat constant ∼ 90 MPa among the native and PEG-crosslinked VLPs. This observation can potentially be explained as follows. In nanoindentation experiments, the shell compression and bending moduli contribute to an effective elastic modulus *E* (*52–54*). Tethered PEG molecules disfavor bending because it results in chain stretching, therefore increasing the bending modulus. Reduction in coat protein positive charges after PEG conjugation reduces the inter-atomic repulsion during compression, therefore decreasing the compression modulus. The higher stiffness of PEG-crosslinked VLPs compared to native VLP is therefore associated with an increase in the shell thickness, as well as stretching the PEG molecules (*55*). The effect of chain stretching became evident where the PEGylated but not crosslinked VLP, namely mPEG_25_-AP205 VLP, was investigated. This VLP showed a lower elastic modulus and stiffness compared to its PEG-crosslinked counterpart, namely bPEG_25_-AP205 VLP (**Figure 3 and Figure 4**).

**Figure 4:**
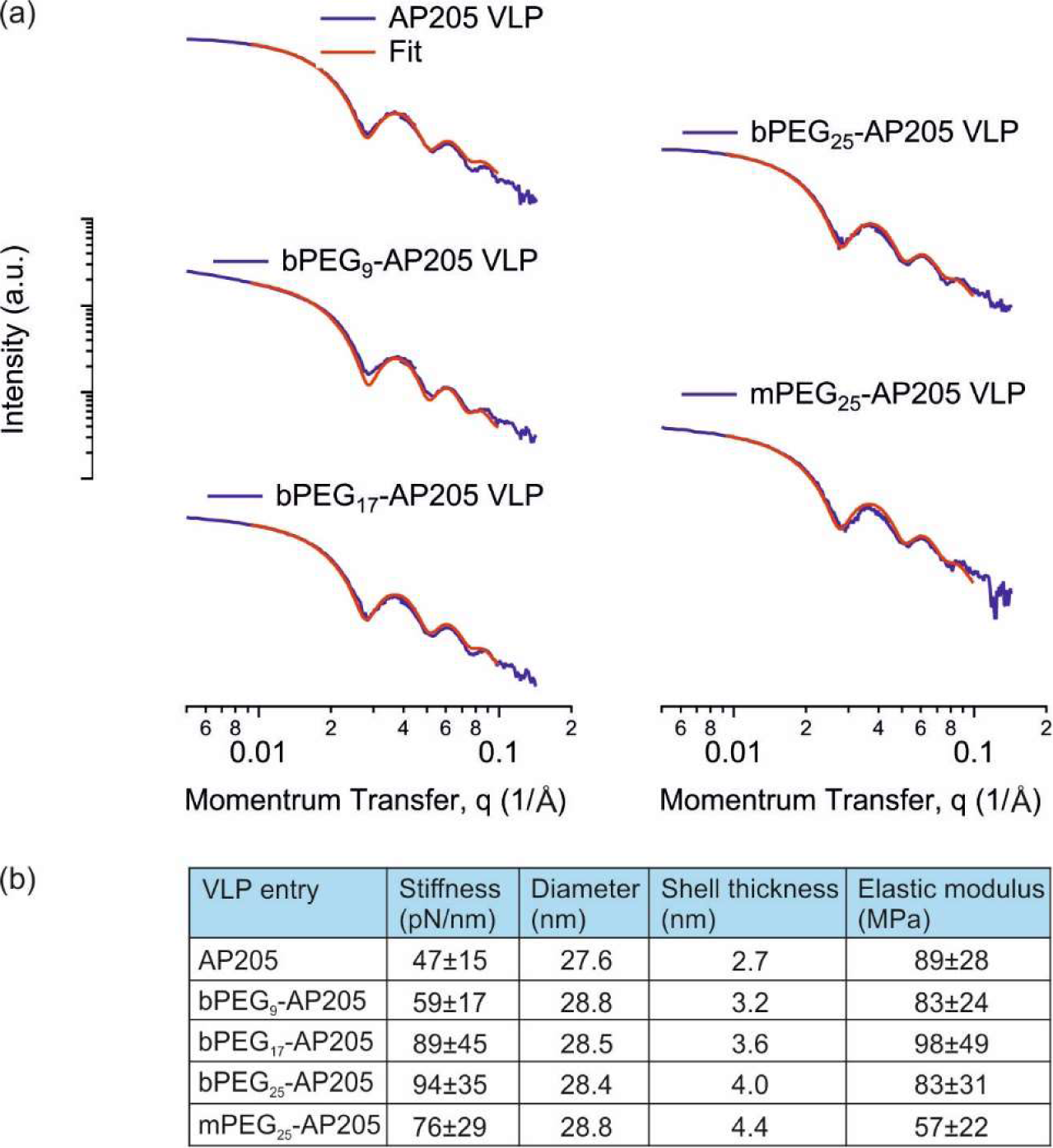
Geometrical and mechanical parameters of native and PEGylated AP205 VLPs. (a) Small angle X-ray scattering (SAXS) spectra of AP205, bPEG_9_-AP205, bPEG_17_-AP205, bPEG_25_-AP205 and mPEG_25_-AP205 VLPs, together with fits to spherical core-shell form factor. (b) Summary panel of the geometrical and mechanical properties of the VLPs.

### Surface-crosslinking does not restrict mucus translocation of AP205 VLP

A major feature of intestinal and respiratory sites, where VLP application is relevant, is the production of mucus. Mucus is an adhesive and viscoelastic gel that covers the epithelium and functions to trap and dispose of foreign objects. In the airways, mucus is constantly pushed out via mucociliary clearance (*56,57*). We used *in vitro* 3D human nasal epithelial tissues to investigate the effect of PEG-crosslinking on translocation across flowing mucus. The tissues were characterized as having fully differentiated mucus producing goblet cells, ciliated cells with motile cilia, as well as other cells of normal human nasal epithelial tissue. The schematic of the experiments is shown in **Figure 5(a)**. On delivery of fluorescently labelled VLPs from the apical side, the VLPs translocated through mucus by diffusion and simultaneously transported in flowing mucus in the direction of mucociliary clearance to edges of the tissue. The lateral distribution and vertical translocation were monitored using laser scanning confocal microscopy (LSCM). In **Figure 5(b)**, representative examples of mean fluorescence intensity (MFI) variation with height collected in the center, [0, 0] mm, prior to VLP delivery (VLP-) and after the delivery of mPEG_25_-AP205 VLP^GelRed^ (VLP+) are shown. In the former case, the location of maximum MFI was at the height level of the semipermeable membrane which was assigned to 0 µm. After VLP delivery, the maximum of MFI shifted to a higher height value (about 40 µm) while the peak associated with semipermeable membrane was observable. The MFI vs height spectra were normalized (norm. MFI) by the MFI values at height = 0 µm, **Figure 5(b)**. The normalized MFI spectrum of VLP-was then subtracted from that of VLP+ to give the net MFI spectrum. The net MFI spectra from the center, [0, 0] mm, together with spectra at positions [−1, 0] mm, [+1, 0] mm, [0, −1] mm and [0, +1] mm, are shown in **Figure 5(c)**. These spectra provided information about the height location of VLP deposition and the net value of MFI at those deposition heights, which corresponded to the translocated amount of VLPs. In all experiments, it was ensured via bright field microscopy that the cilia were beating before and after VLP delivery using stablished procedures (*58*).

**Figure 5:**
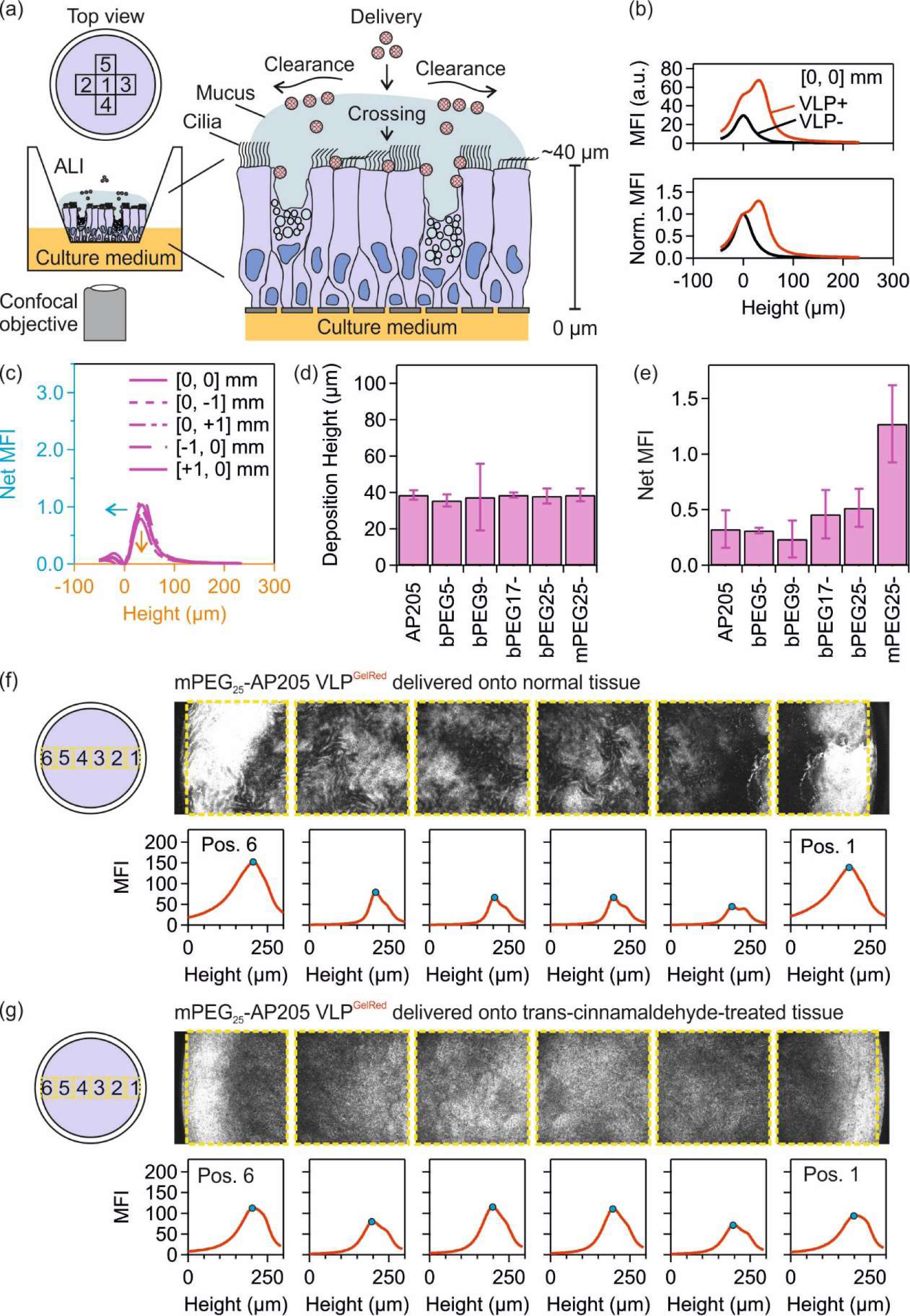
Translocation through mucus of AP205 VLPs in *in vitro* 3D human nasal epithelial tissues. (a) Schematic of *in vitro* nasal epithelial tissue and delivery of fluorescently-labelled virus-like particles (VLPs) from the apical side. Using laser scanning confocal microscopy (objective lens 10x, N.A. 0.3), the lateral distribution and vertical translocation through mucus of VLPs were monitored at several position 1: [0, 0] mm, 2: [−1, 0] mm, 3: [+1, 0] mm, 4: [0, −1] mm and 5: [0, +1] mm. (b) Mean fluorescence intensity (MFI) and normalized MFI (norm. MFI) variation with height collected in the center of tissue 3. MFI and norm. MFI spectra prior to VLP delivery, VLP-, and after the delivery of mPEG_25_-AP205 VLP^GelRed^, VLP+, are shown. The height of semipermeable membrane is set to 0 µm. (c) Subtracted (or net) MFI spectra on five positions showing the height location of VLP deposition and the net MFI. (d, e) The deposition height (d) and the net MFI corresponding to the translocated amount of VLPs (c) of native and PEGylated VLPs on tissue 3. (f) Delivery of mPEG_25_-AP205 VLP^GelRed^ to tissue 3. (g) Delivery of mPEG_25_-AP205 VLP^GelRed^ to trans-cinnamaldehyde-treated tissue 3. The yellow box denotes the area of mean fluorescence intensity (MFI) calculation and blue circle the height location of VLP deposition. In these figures, height starts from above the air-liquid interface (ALI) and increases in the direction of cell layer to below the semipermeable membrane. The height of the semipermeable membrane is approximatively 250 µm. The individual confocal images on lateral positions 1 to 6 were taken at a fixed height distance from the semipermeable membrane.

A typical deposition height and translocated amount of native and PEG-crosslinked VLPs are shown in **Figure 5(d) and (e)**. A similar deposition height between these VLPs was expected as tissues from the same donor (tissue 3) were used. Furthermore, as shown in **Figure 5(e)**, similar translocated amounts between the native and PEG-crosslinked VLPs were obtained. This observation indicated that PEG-crosslinking did not restrict nor increased VLP translocation through mucus. These results were consistent on other tissues (**S8**). We then investigated the translocation behavior of PEGylated but not crosslinked VLP, namely mPEG_25_-AP205 VLP^GelRed^. We found that mPEG_25_-AP205 VLP^GelRed^ showed a similar deposition height (**Figure 5(d)**), however it had an increased translocation amount which exceeded native VLP by two folds (**Figure 5(e)**). Particularly as compared to bPEG_25_-AP205 VLP, this result indicated a higher mucus permeability of the mPEG_25_-AP205 VLP construct. Consistent with previous reports (*59–61*), a potential explanation is an effective steric hinderance interactions with mucin glycoproteins from one-end conjugated PEGs.

We then investigated the effect of outward-swirling mucociliary clearance on the deposition of VLPs across the tissue (*62*). mPEG_25_-AP205 VLP^GelRed^ was delivered in normal tissue condition or when the tissue was treated with trans-cinnamaldehyde. Analysis of the lateral distribution of deposited VLPs across the tissue (positions 1 to 6) under normal conditions (**Figure 5(f)**) showed that a noticeable portion of VLP was cleared to the edge of the plastic insert in flowing mucus while a portion translocated similar to the observations already discussed. However, the lateral distribution of deposited VLPs was more uniform across the tissue when ciliary beating was stopped (**Figure 5(g)**). Additional examples of the deposited VLP distribution when ciliary beating was inhibited are shown in **SI 9**.

### Compatibility of PEG-crosslinking with vaccine function

A major application of VLPs is in mucosal delivery of vaccine antigens. We therefore investigated if PEG-crosslinking may interfere with this function by measuring antibody induction against coat protein in native and PEG-crosslinked VLPs in a subcutaneous vaccination model using C57BL/6J specific pathogen free (SPF) mice, i.e. a setting where vaccine stability is thought not to be a major limitation on performance. Accordingly, subcutaneous injections of VLPs without additional adjuvant were performed on Day 0 and 10. The mice were sacrificed on Day 20 and anti-AP205 serum IgG titers detected using enzyme-linked immunosorbent assay (ELISA). The serum antibody titers against AP205 coat protein are shown in **Figure 6(a)**. As expected, native VLP was immunogenic, and we easily detected specific serum IgG antibodies against AP205 coat protein (*39,40*). Surprisingly, PEG-crosslinking, or simple PEGylation of VLP did not shield immune activation against the coat protein; rather, specific antibody induction was increased. Potentially several mechanisms could be involved, and one cannot make a direct correlation with stiffness. Antibody induction against the PEG coating was also investigated. Consistent with the strong adjuvanticity of VLPs (*63*), the anti-PEG ELISA in **Figure 6(b)** showed that the injection of PEGylated VLPs, compared to the negative control, i.e., native VLP, resulted in anti-PEG IgG induction. Furthermore, the anti-PEG IgG titers seemed to increase with the size of two-end conjugated PEG crosslinker, i.e., to be higher in the order bPEG_9_-AP205 VLP < bPEG_17_-AP205 VLP < bPEG_25_-AP205 VLP.

**Figure 6:**
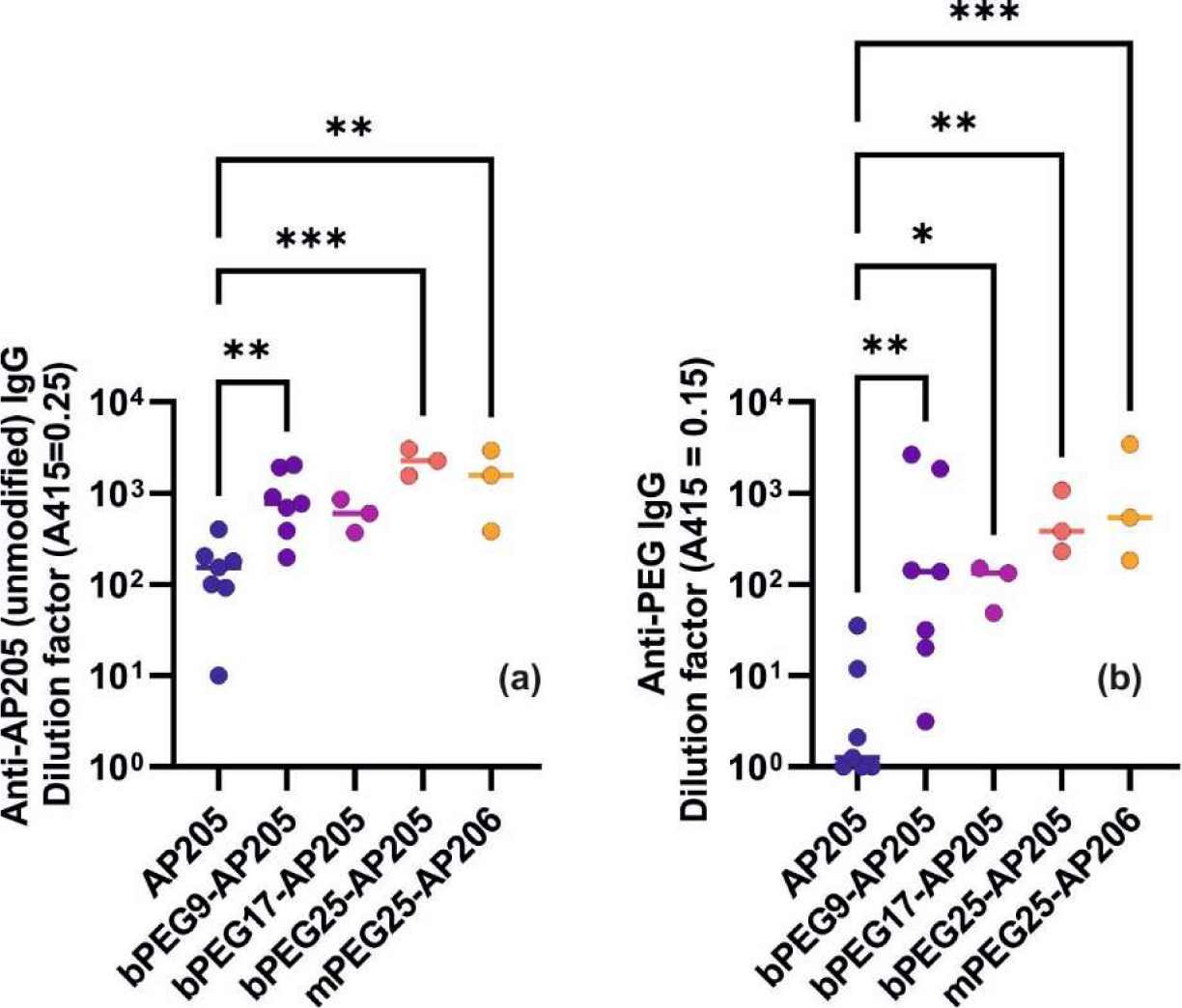
Immune activation by PEG-crosslinked AP205 VLPs. C57BL/6J SPF mice were subcutaneously injected with native (n = 7), PEG-crosslinked (n = 13), or PEGylated but not crosslinked (n = 3) AP205 VLPs on Day 0 and Day 10 and sacrificed on Day 20. Serum antibody (IgG) titers against (a) AP205 coat protein, and (b) PEG was measured by ELISA. Statistical significance was determined by ordinary one-way ANOVA with Dunnett’s multiple comparisons test, with a single pooled variance on log-normalized data (*P = 0.0332; ***P = 0.0021; ***P = 0.0002).

## Discussion

VLPs have similar physical characteristics to their viruses of origin, including size, proteinaceous core-shell structure, and patchy surface charge distribution, and like viruses are prone to aggregation, disintegration, and sticking to surfaces rendering them non-functional (*64*). Enhancing the stability of VLPs is therefore justified in future developments of VLP biotechnology, particularly for mucosal applications. We investigated surface crosslinking as a means to achieve the latter. We found that it increased the colloidal stability of VLPs at low pH (**S6**) as it protected them against proteases. Specifically, while the native AP205 VLP disintegrated in pig gastric fluid at pH 3.0 and 4.7 in less than 5 min, the surface-crosslinked VLPs presented a longer lifetime in the same fluids (**Figure 1** and **S7**). Susceptibility to mechanical agitation was introduced as another limiting factor to VLP mucosal application. Using different force probing techniques, the endocytosis “pulling” force exerted by a cell onto a virus, B cell antigen extraction, and the force of cilia agitation were measured to vary between a few to hundreds of pN (*44–50*). Although this force range is generally below the yield force values of VLPs, when applied repeatedly as they do *in vivo*, they can disintegrate the capsids due to fatigue (*16-18, 22-25*). Therefore, it is also logical to mechanically reinforce VLPs and surface-crosslinking was found to increase the stiffness and strength of AP205 VLP (**Figure 2 and Figure 4**).

Mucosal applications of VLPs are faced with other physiological challenges, namely translocation through a thick mucus layer. Particularly in the airways, the mucus is subject to mucociliary clearance which further reduces VLP translocation efficiency by pushing the mucus out of the airways (*56*). Experiments using *in vitro* 3D human nasal epithelial tissues showed an effective translocation of native and surface-crosslinked AP205 VLPs, while mucociliary clearance transported mucus at a speed of about 40–200 µm/s (measured at 1–3 mm from the tissue center, (*62,65*)) in a swirling rotation to the edges of the tissue (**Figure 5**). Is mucus a poor barrier to VLPs of a size below about 100 nm, including AP205 VLP (27.6 nm), and VLPs of human papilloma virus and Norwalk virus (respectively, 55 nm and 38 nm, (*66*))? Considering viruses that infect mucosal surfaces, including polio (28 nm), hepatitis B (43 nm), adenoviruses (60–90 nm), rotavirus (75 nm), human immunodeficiency virus (120 nm) and herpes simplex virus (180 nm) (*67*), one may conclude that small viruses are capable of deeply penetrating the mucus. This is expected when the pore size of mucus gel (≥ 50 nm) exceeds the size of these viruses. However, penetration is further facilitated by the patchy surface charge distribution on capsids (*66,67*). Therefore, structural point mutations as well as macromolecule entanglement with mucin glycoproteins can alter the diffusion of VLPs in mucus (*68*).

Finally, our results suggest that the material properties of VLPs are relevant variables that should be further investigated to improve mucosal applications. As schematically shown in **Figure 1(a)**, both chemical, enzymatic, and physical stresses can affect the efficacy of VLP delivery to the respiratory or intestinal mucosal epithelium. In this work, crosslinking of multiple surface sites was shown to be a promising approach to improve and extend the range of VLP mucosal functions by increasing resistance to degradation and improving physical stability. These modifications actually slightly enhance even parenteral vaccination indicating that the tethers did not shield the coat proteins from immune recognition, and that increase biochemical stability may also be beneficial systemically (**Figure 6**). PEG immunogenicity is currently an unwanted side-effect that may be overcome with use of alternative crosslinkers in the future. Critically, our work demonstrates the potential to improve the current VLP system to generate structures compatible with overall enhanced functionality, strength, stability and immunogenicity at mucosal surfaces.

## Materials and Methods

### Experimental Design

Surface crosslinking of virus-like particle (VLP) capsids using polyethylene glycol (PEG) tethers was investigated against physiological challenges to assess its performance on mucosal surfaces. The model native VLP was derived from the coat protein of Acinetobacter phage, namely AP205 VLP. PEG tethers included two-end functional PEG molecules of various molecular weights. The PEG-crosslinked AP205 VLPs were initially evaluated in terms of the reaction products and the sites of PEG conjugation to reactive surface sites (using a combination of dynamic light scattering, transmission electron microscopy, reducing SDS-polyacrylamide gel electrophoresis, electrospray ionization mass spectroscopy and nano ultra-performance liquid chromatography-MS) and then investigated for the aggregation behavior against pH (using dynamic light scattering), disintegration against digestive enzymes from pig and mouse gastric fluid (using agarose gel electrophoresis), resistance to mechanical force (using atomic force microscopy), and penetration in mucus in the presence of motile cilia and mucociliary clearance (using *in vitro* 3D human nasal tissues).

### Production of AP205 VLP

The biosynthesis of AP205 VLP is reported in details in our previous study (*25*). Briefly, it includes the expression of recombinant AP205 coat proteins in bacterial cytoplasm which then self-assemble into AP205 VLPs. After the expression of VLP, a series of steps consisting of bacterial lysis, digestion of RNA and DNA contaminants, and extraction of lipids were performed prior to precipitation and dialysis against PBS to produce purified AP205 VLPs. Plasmids were generated by custom DNA synthesis (Twist Bioscience, San Francisco, CA, USA) of the expression cassettes and cloning into the pRSFDuet-1 (Novagen) backbone between the PfoI and BshTI restriction sites. Sequences of the plasmids are available via the ETH-Zürich research collection at https://doi.org/10.3929/ethz-b-000556182. The sequence of AP205 monomer and the description of plasmid are reported in **S1**.

### PEGylation of AP205 VLP

The functional group on the extremities of homobifunctional PEG crosslinker molecules (bPEG_n_, n = 5 to 25) and on one extremity of monofunctional PEG molecules (mPEG_n_, n = 12 and 25) was NHS-ester which is reactive to primary amine of lysine residues and N-terminus of AP205 coat proteins. Reactions with native AP205 VLP proceeded at 1–100x molar ratio of crosslinker molecule to AP205 monomer (molar ratio = [functional moleucle] [AP205 CP]). The PEG molecule was dissolved in anhydrous and molecular sieves dried dimethyl sulfoxide (DMSO, Sigma-Aldrich) to a concentration of 100 mM. An appropriate amount was then added to AP205 VLP suspension at 4 mg/ml in PBS. The reaction was left for 24 h at 4°C. Then after, it was quenched by adding Tris•HCl (Trizma^®^ base, Sigma-Aldrich, and hydrochloric acid 37%, VWR) at 20–50 mM and incubating for 15 min. Unreacted PEG molecules and Tris were removed using Zeba Spin Desalting Column (ThermoFisher) with molecular weight cut-off at 7 kDa. Other reactive molecules including BS^3^ and Sulfo-NHS were used similarly; however, they were initially dissolved to 100 mM in PBS. A summary of the physical properties of these molecules is provided in **S1**.

### Fluorescent staining of VLPs

We defined a mixing ratio *x* = mass of stain⁄mass of VLP = 10^-2^, and a final stained VLP concentration equal to 5.3 mg/ml. (To calculate the mass of stain, we assumed 10,000x to correspond to 10 mg/ml.) GelRed^®^ (10,000x in water, biotium) or SYBRGold^TM^ (10,000x in DMSO) were used and staining proceeded for 30 min. To remove unbound dye, the solution of stained VLP was buffer exchanged with pure PBS using Zeba™ Spin Desalting Columns (7,000 MWCO, ThermoFisher). We note that since the columns contained sodium azide, it was important to buffer the column with pure PBS prior to cleaning the stained VLP solution. The stained VLP solution was finally filtered using 0.2 µm centrifuge filters (Corning^®^ Costar^®^ Spin-X^®^ centrifuge tube filters) to remove possible aggregates. The intensity of stained VLPs was monitored over time using laser scanning confocal microscopy and showed no noticeable reduction over a course of at least two weeks.

### Transmission electron microscopy (TEM)

4 µl VLP solution (0.1 mg/ml) was deposited on glow-discharged (45 sec, 25 mA, negative charge) carbon-coated grids (Ted Pella) for 1-5 min, after which the excess liquid was removed using filter paper. Then the grid was rinsed with 5 µl of PBS to remove loosely bound VLPs. Subsequently, 3 μl of 2% uranyl acetate was applied and immediately removed with filter paper prior to a second treatment with 3 μl of 2% uranyl acetate for 20 sec. Excess liquid was again collected using filter paper and the grid was let dry at room temperature (RT). Data acquisition was performed using a Morgani transmission electron microscope operating at 100 kV acceleration voltage.

### Electrospray ionization mass spectrometry (ESI-MS)

ESI-MS was acquired on a Synapt G2-Si quadrupole time-of-flight spectrometer (Waters, UK) with a capillary voltage of 3 V, a cone voltage of 50 V and a source temperature of 100°C. VLPs at a concentration of 1 mg/ml were reduced by dithiothreitol (DTT) at a final concentration of 50 mM. The reduction was performed for 1 h at RT and at pH 8. The samples were acidified with 1% formic acid (Thermo, USA), desalted using C<u>4</u> Zip Tips (Millipore, USA) and analyzed in methanol:2-propanol:0.2 % formic acid (30:20:50). The solutions were infused through a fused silica capillary (ID 75 µm) at a flow rate of 1 μl/min and sprayed through Pico Tips (ID 30 µm). The last were obtained from New Objective (Woburn, MA, USA). Recorded m/z data were deconvoluted using the MaxEnt1 software (Waters, UK) with a resolution of the output mass of 0.5 Da per channel and a Uniform Gaussian Damage Model at the half-height of 0.5 Da.

### Nano ultra-performance liquid chromatography-MS (nanoUPLC-MS/MS)

NanoUPLC-MS/MS was acquired on Thermo FUSION mass spectrometer (Thermo, Germany) coupled to a nano-M-class UPLC system (Waters). BPEG_9_-AP205 VLP at a concentration of 1 mg/ml was digested by incubating 3 µl of the sample with 42 µl of 25 mM ammonium bicarbonate (pH 8.5) and 1 µl of TCEP (100 mM) for 1 h at 37°C. Reduced protein was incubated with 2 µl chloroacetamide (500 mM) for 1 h at 37°C. Then after, 5 µl trypsin (0.1 µg/µl in 10 mM HCl) was added and the solution further incubated overnight at 37°C. When needed the pH was adjusted to pH 8. At the end of this procedure, the sample was dried using SpeedVac. Digested and dried sample was dissolved in 20 µl ddH_2_O with 0.1% formic acid and diluted 5 times before transferring to autosampler vials. Peptides were resuspended in 2.5% acetonitrile with 0.1% formic acid and loaded onto a nanoEase M/Z Symmetry C18 (75 μm × 20 mm, 100 Å, 3 μm particle size) and separated on a nanoEase M/Z HSS C18 T3 (75 μm × 150 mm, 130 Å, 1.7 μm particle size) at a constant flow rate of 300 nl/min, with a column temperature of 50°C and a linear gradient of 2−32% acetonitrile/0.1% formic acid in 79 min, and then 32-45% acetonitrile/0.1% formic acid in 10 min, followed by a sharp increase to 98% acetonitrile in 2 min and then held at 98% for another 10 min. Mass spectrometer was operated under data-dependent acquisition (DDA), one scan cycle comprised of a full scan MS survey spectrum, followed by up to 12 sequential higher-collisional energy (HCD) MS/MS on the most intense signals above a threshold of 1e4. Full-scan MS spectra (600–2000 m/z) were acquired in the FT-Orbitrap at a resolution of 70,000 at 400 m/z, while HCD MS/MS spectra were recorded in the FT-Orbitrap at a resolution of 35,000 at 400 m/z. HCD was performed with a target value of 1e5 and normalization collision energy 25 NCE (normalize collisional energy) was applied. Auto-gain control (AGC) target values were 5e5 for full Fourier transform (FT) MS. For all experiments, dynamic exclusion was used with a single repeat count, 15 s repeat duration, and 30 s exclusion duration. The acquired MS data were processed for identification using the Byonic search engine (PMI). The spectra were searched against AP205 VLP sequence and AP205 VLP sequence with E. *Coli* database. The following modifications were included: (a) variable modifications, namely oxidation at N-terminus methionine and carbamidomethylation at cysteine (C), and (b) user-defined modifications, namely 478.2565 (C_22_O_11_H_38_) was set for PEG_9_-crosslinked peptides and 496.2565 (C_22_O_12_H_40_) for one-end PEG_9_ conjugation at a lysine residue or N-Terminus. In this analysis, the raw data were analyzed by two software, PMI Byonic search engine and SIM 1.5.5.3 (Spectrum identification machine) (*69*). In addition, two FastA databases with and without starting methionine were used to confirm the N-terminus.

### SDS-polyacrylamide gel electrophoresis (SDS-PAGE)

Denaturing and reducing SDS-PAGE was carried out in 13% Tricine gels. Samples of 3 μg of VLP were denatured and reduced in Laemmli sample buffer at 95°C for 5 min prior to electrophoresis. Proteins were visualized by Coomassie staining.

### Dynamic light scattering (DLS)

DLS was performed at a fixed angle of 173° by averaging 3 runs of 30-s long each using Zetasizer Nano (Malvern Panalytical). The time correlation function of scattered intensity was analyzed by the cumulant as well as CONTIN methods. The VLP concentration was initially adjusted to 0.1 mg/ml in filtered PBS prior to measurements.

### Gastric fluid collection

Pig gastric content was collected from the stomach of conventionally reared, outbred pigs with different genetic contributions of French-Swiss Landrace and Swiss Large White. Total gastric content was collected. Mouse gastric content was obtained by collecting total stomach content from several specific opportunistic pathogen-free (SPF) C57BL/6J WT mice. The gastric content from several mice was pooled to have sufficient volume. Pig or mouse gastric content was centrifuged for 5 min at 16000 rpm to separate the non-digested food particles. After the collection of supernatant (gastric fluid), it was filtered using 0.22 µm sterile filter and the pH measured using a laboratory pH-meter.

### Agarose gel electrophoresis

VLPs from 1 mg/ml preparation were diluted 1:10 in pig or mouse gastric fluid and incubated for 5 min, 30 min, 1 h and 1.5 or 2 h at 37°C. After each incubation, the samples were immediately placed on ice. Native agarose gel electrophoresis of PBS- or gastric fluid-treated VLPs was carried out in 0.6% agarose in TAE buffer (40 mM Tris, 0.1% acetic acid, 1 mM EDTA) with 0.5 μg/ml of ethidium bromide for visualization. Samples of 5 μg VLP were electrophoresed at 120 Volts for 1 h, then imaged on a UV transilluminator.

### Atomic force microscopy (AFM)

The schematic of AFM measurements is shown in **Figure 2(a)**. After an initial calibration, which also included the calibration of PeakForce tapping mode, on freshly cleaved mica, the tip was brought in proximity to VLPs immobilized at a concentration of 5 µg/ml on Highly Oriented Pyrolytic Graphite (HOPG, Ted Pella, Inc.) The calibrated parameters included stiffness (59.4–82.0 pN/nm) using thermal method and optical lever sensitivity using cantilever–mica contact region (*70*), and drive3 amplitude sensitivity. During the transport of cantilever from mica to HOPG, the cantilever remained wet, and the laser spot was checked to remain on the same spot. The calibration values were re-measured on mica at the end of the measurements to detect any noticeable variation to the calibrated parameters. Single VLPs were localized on HOPG by imaging in PeakForce tapping mode (peak force 50 pN, tapping frequency 1 kHz, amplitude 30 nm). Generally, images of individual VLPs were acquired at a scan rate of 0.7 Hz with 200×200 or 500×500 nm^2^ scan size and a resolution of 64 points per line, leading to a pixel size of 3 nm. After localization, we used “Point & Shoot” function (NanoScope, Bruker) to place the AFM tip on the apex of selected VLPs before applying indentation using Ramp mode. The VLPs were indented up to a threshold force value of 400 pN at a preset frequency of 1 Hz, which resulted in an indentation force rate of about (4–9)×10^3^ pN/s. Indented particles were imaged again in PeakForce tapping to detect signs of degradation, drift, or displacement due to tip lateral forces. The height of individual VLPs before (*H_i_*) and after (*H _f_*) indentation were calculated by placing a trace line over the particles on images previously flattened using Gwyddion 2.47 (*71*). For stiffness evaluation, we proceeded as follows: deflection versus piezo displacement was converted to force versus tip separation using protocols written in Igor Pro (Wavemetrics). Fit to force versus indentation distance (*F* − *δ*) curve in the linear response part of the tip–VLP contact region gave the stiffness (*k* = *dF/dδ*) and the indentation (*δ* = Δ*F/k*). The fit was performed in the force range 50 pN ≤ *F* ≤ 250 pN, and only those VLPs with indentation range 2.0 ≤ *δ* ≤ 6.0 nm were selected for averaging. Only one indentation per VLP was used for the evaluations. The indentation force rate (*Ḟ*) was calculated from a linear fit to the force versus indentation time (*F* − *t*) curve in the same region were the stiffness of VLP was calculated. Strain rate was then calculated from the relation 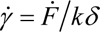. Average stiffness was calculated from at least duplicate independent VLP preparations. UV-ozone (Novascan) cleaned AC40 BioLever Mini cantilevers (Bruker) were used. Dimension FastScan (Bruker) was used for imaging and force spectroscopy.

### Small angle X-ray scattering (SAXS)

We used a Xeuss 2.0 (Xenocs, France) instrument with a microfocused X-ray source at the Laboratory Léon Brillouin (LLB NIMBE CEA Saclay). The Cu Kα radiation (λ_Cu Kα_ = 1.5418 Å), and the data were collected by a 2D Pilatus 1 M detector (Dectris, Switzerland). The scattering vector *q* = 4*π/λ* sin(*θ* 2), with *θ* being the scattering angle was calibrated using silver behenate. Data were collected and azimuthally averaged to produce unidimensional intensity versus *q*, with *q* interval of 0.005 to 0.1 Å^-1^. The measurements were performed at 25°C. The scattering intensity was collected for 2600 s. The intensity was corrected for sample thickness, transmission, and acquisition time. The modelling of the integrated data was performed with a core-shell form factor, and a hard sphere structure factor, when needed, using the SasView software.

### *In vitro* 3D human nasal epithelial tissue

*In vitro* human nasal epithelial tissues from single donors (MucilAir^TM^) were obtained from Epithelix (Switzerland). The tissues were maintained according to company’s protocols and included tissue 1 from a Caucasian 64-year-old female subject (MD0860), tissue 2 from a Caucasian 46-year-old male subject (MD0871) and tissue 3 from a Caucasian 38-year-old male subject (MD0774). The subjects were non-smokers and had no history of respiratory pathology.

### Inhibition of mucociliary clearance

Ciliary beating was inhibited using trans-cinnamaldehyde (97%, C80687, Sigma-Aldrich). Trans-cinnamaldehyde was dissolved in MucilAir^TM^ culture medium to a concentration of 30 mM. To improve solubility, freshly prepared solution was sonicated for 15 min and the solution was kept at 37°C prior to filling the basal channel of MucilAir^TM^ monodonor nasal epithelial tissue. After 20 min of incubation, the ciliary beating stopped, and no recovery was observed for up to 2 h (*72*).

### Delivery of VLPs to *in vitro* 3D human nasal epithelial tissue

Tissues were incubated in a humidified-CO_2_ chamber at 37°C. After a period of equilibration 0.5–1 h, 5 µl of VLP solution (5.3 mg/ml) was delivered from the apical side. The delivery was onto the center of the tissue.

### Laser scanning confocal microscopy (LSCM)

Lateral distribution and vertical translocation of fluorescently labelled VLPs through mucus were then monitored using laser scanning confocal microscopy (LSCM, Leica SP5 confocal microscope). We used a 10x objective lens with N.A. 0.3. Excitation wavelength *λ*_ex_ = 488 nm was used for SYBRGold^TM^-stained VLPs and Salmonella vaccine with detection wavelength *λ*_em_ = 500 − 590 nm. *λ*_ex_ = 514 nm was used for GelRed^®^-stained VLPs with *λ*_em_ = 550 − 650 nm. The channel spectral detection was set to HyD (Gain equal to 100). The mean fluorescence intensity (MFI) variation with height was collected in the center [0, 0] mm, and on positions [−1, 0] mm, [+1, 0] mm, [0, −1] mm and [0, +1] mm prior to, during and after VLP delivery. Leica LAS AF software was used for saving the images. Analysis was performed using ImageJ.

### Mice

All animal experiments were performed in accordance with Swiss Federal regulations approved by the Commission for Animal Experimentation of the Kanton Zurich (license ZH009/2021, Kantonales Veterinäramt Zürich, Switzerland). SPF C57BL/6J WT mice were used in all experiments. Mice were bred and housed in individually ventilated cages with a 12 h light/dark cycle in the ETH Phenomics Center (EPIC, RCHCI), ETH Zürich, and were fed a standard chow diet. All mice included in experiments were 7 weeks or older and objectively healthy as determined by routine health checks. Wherever possible an equal number of males and females was used in each experimental group.

### Subcutaneous injection

50 µg of VLP in 100 µl of PBS was subcutaneously injected into the loose skin over the interscapular area of C57BL/6J SPF mice on Day 0 and Day 10. The mice were euthanized on Day 20. Blood was collected by cardiac puncture, and serum separated by spinning 1.1 ml serum gel tubes (Sarstedt) at 10,000 g for 5 min at 20°C in a conventional table-top centrifuge, and heat-inactivated for 30 min at 56°C.

### Enzyme-linked immunosorbent assay (ELISA) for detection of anti-AP205 IgG response

50 µl of AP205 VLP solution (5 µg/ml) was pipetted into an appropriate number of wells of ELISA plates (4 Nunc MaxiSorp flat bottom 96-well plates) and the VLP allowed to adsorb overnight at 4°C. The wells were washed three times with wash buffer (0.05% Tween 20 in PBS) and blotted. The wells were blocked using 150 µl of blocking buffer (2% bovine serum albumin in PBS) for 1.5 h at RT. Subsequently, the wells were washed five times with wash buffer and blotted. Mouse sera were diluted ten times in blocking buffer, and sequentially in 1:3 dilution steps added to the wells of ELISA plates. The plates were incubated for 2 h at RT to allow the binding of serum IgG to AP205 proteins. Goat anti-mouse IgG-HRP (Sigma) was diluted a thousand-fold in blocking buffer, and 50 µl of solution added to the wells and incubated for 1 h at RT. The wells were washed five times using wash buffer and blotted. Finally, 150 µl of freshly prepared substrate solution (consisting of 0.1 M NaH_2_PO_4_ (substrate buffer), 2,2’-azino-bis(3-ethylbenzothiazoline-6-sulfonic acid (1–2 mg per 10 ml of substrate buffer), and H_2_O_2_ (10 µl per 10 ml)) was added to each well. The plate was incubated in the dark for 40 min. On Tecan (Tecan Infinite 200 PRO microplate reader), absorbance was measured at 415 nm. The reference wavelength for the subtraction of background signal was set to 480 nm.

### Enzyme-linked immunosorbent assay (ELISA) for detection of anti-PEG IgG response

Mouse sera were diluted ten times in PEG dilution buffer (PEGD50-1, Life Diagnostics, Inc.), and sequentially in 1:3 dilution steps added to the wells of PEG-BSA coated ELISA plates (PBSA20PL, Life Diagnostics, Inc.) The plates were incubated for 2 h at RT to allow antibody binding. The wells were washed five times with 300 µl PEG wash buffer (PEGW50-20, Life Diagnostics, Inc.) and blotted. Goat anti-mouse IgG-HRP (AP124P, Sigma-Aldrich) was diluted a thousand-fold in PEG dilution buffer and 100 µl of this solution was added to each well with an incubation of 1 h at RT. The wells were washed and blotted again. Finally, 150 µl of freshly prepared substrate solution was added to each well. The plates were incubated in the dark for 40 min. On Tecan (Tecan Infinite 200 PRO microplate reader), absorbance was measured at 415 nm. The reference wavelength for the subtraction of background signal was set to 480 nm.

### Statistical Analysis

AFM and LSCM results were expressed as mean ± standard deviation analyzed by Igor Pro from WaveMetrics. *In vivo* mouse vaccination results were assessed using ordinary one-way ANOVA with Dunnett’s multiple comparisons test using GraphPad Prism (*P = 0.0332; ***P = 0.0021; ***P = 0.0002).

## Acknowledgments

We acknowledge the contributions from Till Germerdonk in DLS measurements, and discussions with Patrick Zueblin. We also acknowledge the time for SAXS measurements provided by the Leon Brillouin Laboratory (LLB), and the support of Fabrice Cousin.

## Funding

European Research Council (ERC) Grant number 865730 (MR, YT, AA and ES)

Swiss National Science Foundation, Switzerland, rant number IZSEZ0_212991 (MR)

Swiss National Science Foundation, Switzerland, grant numbers 40B2-0_180953 (ES)

Swiss National Science Foundation, Switzerland, grant numbers 310030_185128 (ES)

National Center for Competence in Research in Microbiomes, Switzerland (ES)

Botnar Research Centre for Child Health Multi-investigator project “Microbiota Engineering for Child Health” (ES)

Gebert Rüf Microbials, grant number GR073_17 (ES)

Cystic Fibrosis Trust UK SRC 016 (PC, JK, RF)

EU ITN PhyMot (EC)

## Author contributions

Conceptualization: MR

Methodology: MR, PC, RM, ES

Investigation: MR, AA, SG, VL, TK, YT, WLB, EC, SC, CWL, JK

Writing—original draft: MR

Writing—review & editing: All authors

Funding acquisition: MR, PC, ES

## Data and materials availability

All data used in the analyses are available without limitation from the first author.

