## Supplementary Information for "Surface crosslinking of virus-like particles increases resistance to proteases, low pH, and mechanical stress for mucosal applications"

Ahmed Ali *et al.*

*Milad Radiom; Emma Slack

### S1. AP205 coat protein sequence and the molecules used to modify AP205 VLP

#### AP205 sequence

MANKPMQPITSTANKIVWSDPTRLSTTFSASLLRQRVKVGIAELNNVSGQYVSVYKRPAPKPEGCADACVIMPNENQS
IRTVISGSAENLATLKAEWETHKRNVDTLFASGNAGLGFLDPTAAIVSSDTTAGSGGAHATANATAHHHHHH

The plasmid used to produce AP205 monomers in *E.* Coli cytoplasm is reported in the table below.

| Plasmid | Description | Source |
|---|---|---|
| pTK190 | Expression construct for AP205 VLP. pRSFDuet backbone with a custom expression cassette consisting of a T5 promoter with dual lac operators, RBS, open reading frame encoding the AP205 coat protein C-terminally fused to a 6xHis tag, followed by a T7 terminator. | This study |

#### Molecules used to modify AP205 VLP

| Molecule<br>Company, product code | Homobifunctional/<br>monofunctional | Spacer length (nm) | Molecular weight (Da) | Name of conjugated AP205 VLP |
|---|---|---|---|---|
| $BS^3$<br>bis(sulfosuccinimidyl)suberate<br>Thermo Scientific 21580 | Homobifunctional | 1.1 | 572.4 | $BS^3$-AP205 |
| $BS(PEG)_5$<br>Bis(succinimidyl) penta(ethylene glycol)<br>Thermo Scientific 21581 | Homobifunctional | 2.2 | 532.5 | $bPEG_5$-AP205 |
| $BS(PEG)_9$<br>Bis(succinimidyl) nona(ethylene glycol)<br>Thermo Scientific 21582 | Homobifunctional | 3.6 | 708.7 | $bPEG_9$-AP205 |
| $BS(PEG)_{13}$<br>Bis-$dPEG_{13}$-NHS ester<br>Quanta Biodesign 10954 | Homobifunctional | 5.0 | 884.9 | $bPEG_{13}$-AP205 |
| $BS(PEG)_{17}$<br>Bis-$dPEG_{17}$-NHS ester<br>Quanta Biodesign 10979 | Homobifunctional | 6.4 | 1061.1 | $bPEG_{17}$-AP205 |
| $BS(PEG)_{21}$<br>Bis-$dPEG_{21}$-NHS ester<br>Quanta Biodesign 10956 | Homobifunctional | 7.9 | 1237.3 | $bPEG_{21}$-AP205 |
| $BS(PEG)_{25}$<br>Bis-$dPEG_{25}$-NHS ester<br>Quanta Biodesign 10968 | Homobifunctional | 9.3 | 1413.6 | $bPEG_{25}$-AP205 |
| Sulfo-NHS<br>N-hydroxysulfosuccinimide<br>Thermo Scientific 24510 | Monofunctional | 0 | 217.1 | Sulfo-NHS-AP205 |
| m-$dPEG_{12}$-NHS ester<br>Quanta Biodesign 10262 | Monofunctional | 4.5 | 685.8 | $mPEG_{12}$-AP205 |
| m-$dPEG_{25}$-NHS ester<br>Quanta Biodesign: 11291 | Monofunctional | 9.0 | 1258.4 | $mPEG_{25}$-AP205 |

### S2. PEG-crosslinking at varying reaction molar ratios

Experiments using dynamic light scattering helped to find a mixing ratio range wpreventing VLP aggregation. Hereafter, to prevent a large excess of free PEG molecules, all reactions were carried out at 10x molar ratio of crosslinker molecule to AP205 monomer.

##### bPEG$_9$-AP205 VLP

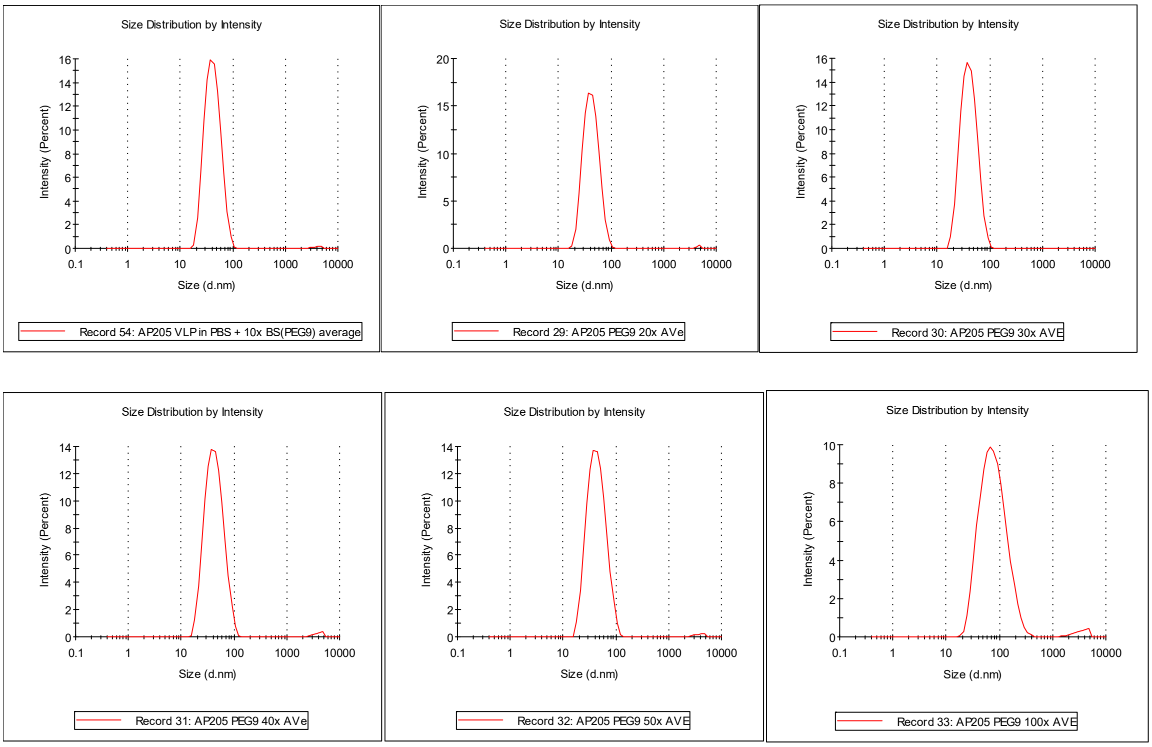

Figure S2-1: Hydrodynamic diameter of bPEG$_9$-AP205 VLP at varying molar ratio of BS(PEG)$_9$ to AP205 monomer.

Table S2-1: Summary of the measurements from Figure S2-1 Cumulant and Contin analysis algorithms were used for data analysis. $D_h$ is hydrodynamic diameter and PDI polydispersity index.

| Molar ratio | Cumulant $D_h$ (nm) | PDI | Contin (1st peak) $D_h$ (nm) |
|---|---|---|---|
| 10x | 35.0 | 0.10 | 38.8 |
| 20x | 40.7 | 0.17 | 43.6 |
| 30x | 37.1 | 0.14 | 42.0 |
| 40x | 37.4 | 0.23 | 44.8 |
| 50x | 40.0 | 0.18 | 45.6 |
| 100x | 67.7 | 0.24 | 86.6 |

#### bPEG$_{25}$-AP205 VLP

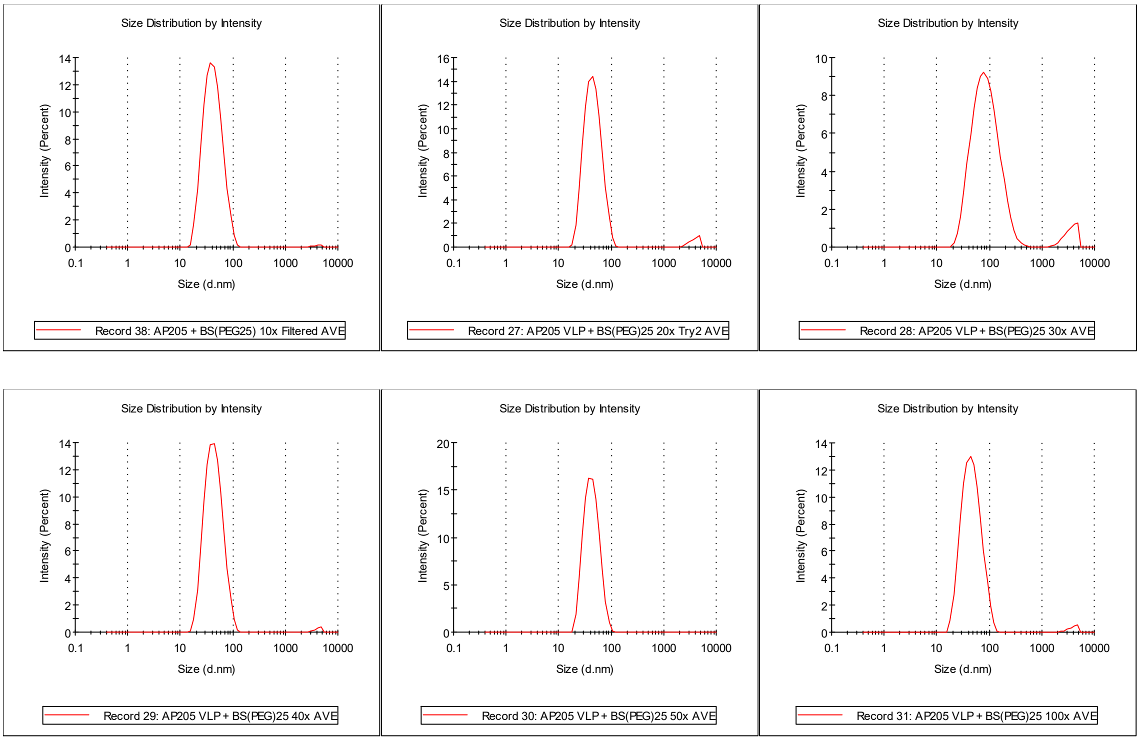

Figure S2-2: Hydrodynamic diameter of bPEG$_{25}$-AP205 VLP at varying molar ratio of BS(PEG)$_{25}$ to AP205 monomer.

Table S2-2: Summary of the measurements from Figure S2-2. Cumulant and Contin analysis algorithms were used for data analysis. $D_\mathrm{h}$ is hydrodynamic diameter and PDI polydispersity index.

| VLP | Cumulant $D_\mathrm{h}$ (nm) | PDI | Contin (1$^{st}$ peak) $D_\mathrm{h}$ (nm) |
|---|---|---|---|
| 10x | 38.8 | 0.16 | 44.2 |
| 20x | 43.8 | 0.21 | 46.9 |
| 30x | 86.0 | 0.21 | 97.6 |
| 40x | 39.7 | 0.19 | 45.7 |
| 50x | 39.8 | 0.13 | 43.9 |
| 100x | 42.9 | 0.20 | 48.9 |

### S3. Morphology of PEGylated VLPs

Using transmission electron microscopy (TEM), we investigated the morphology of PEGylated AP205 VLPs. As expected, the results shown in figure below demonstrated a spherical geometry after PEGylation with BS($PEG_9$), BS($PEG_{13}$), BS($PEG_{17}$), BS($PEG_{21}$) and BS($PEG_{25}$).

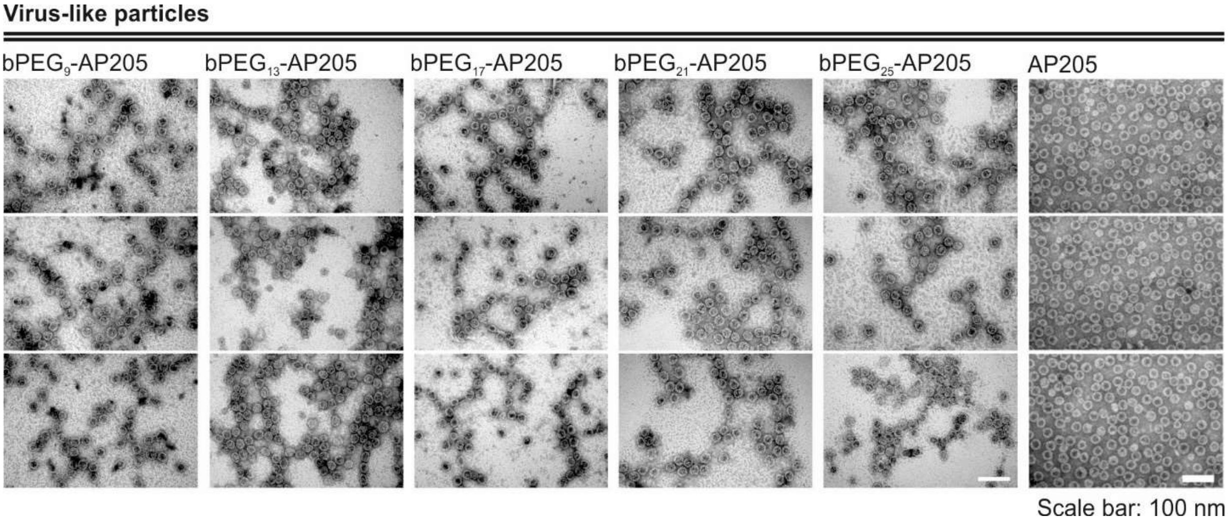

Figure S3-1: TEM images of $bPEG_9$-AP205 VLP, $bPEG_{13}$-AP205 VLP, $bPEG_{17}$-AP205 VLP, $bPEG_{21}$-AP205 VLP, $bPEG_{25}$-AP205 VLP as well as AP205 VLP.

### S4. Mass spectrometric characterization of PEGylated VLPs

#### Electrospray ionization mass spectroscopy (ESI-MS)

From these measurements, the average MW of AP205 monomer was measured to be 15768 Da. Reaction with BS(PEG$_9$) at 1x molar ratio resulted in a significant fraction of AP205 monomers remaining either non-PEGylated or singly-PEGylated. Increasing the molar ratio to 5x decreased the population of non-conjugated AP205 monomers, while peaks at doubly- and triply-PEGylated AP205 monomers appeared. At 10x molar ratio, there was no population of non-PEGylated AP205 monomers, and we found a maximum of four PEG molecules per AP205 monomer. The variation of average MW between the individual peaks corresponded to the MW of one-end conjugated BS(PEG$_9$). The reactions with Bis-dPEG$_{13}$-NHS ester, Bis-dPEG$_{17}$-NHS ester and Bis-dPEG$_{25}$-NHS ester followed a similar behavior; however, at 10x molar ratio, a maximum of three PEG molecules per AP205 monomer was detected.

##### ESI-MS of AP205 VLP

In the figure below, the mass spectrum of AP205 VLP is divided into three regions corresponding to the molecular weight of AP205 monomer (MW 15768 Da), dimer (MW 31536 Da), and trimer (MW 47304) which are indicated on each subfigure (respectively, left, middle and right).

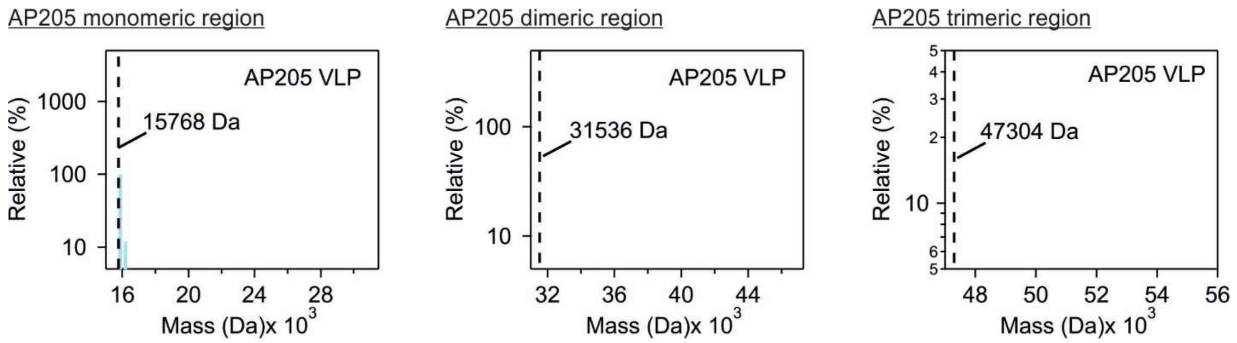

Figure S4-1: ESI-MS data of AP205 VLP.

##### ESI-MS of PEGylated AP205 VLPs

In the figures below, the mass spectra of PEGylated AP205 VLPs are shown. The spectra are divided into three regions corresponding to the molecular weight of AP205 monomer (MW 15768 Da), dimer (MW 31536 Da), and trimer (MW 47304 Da) which are indicated on each subfigure (respectively, left, middle and right). The tables following the figures summarize the occurrence of peaks in the monomer region. The entries, only if they are highlighted in green, show the occurrence of unconjugated AP205 monomer and the occurrence of conjugated AP205 monomers with 1 to 8 PEG molecules. The percentage abundance of each form, relative to the occurrence of the most populated form (= 100%), are indicated below the molecular weight from ESI-MS. Only peaks above 10% occurrence are indicated in the tables.

#### ESI-MS of bPEG$_9$-AP205 VLP

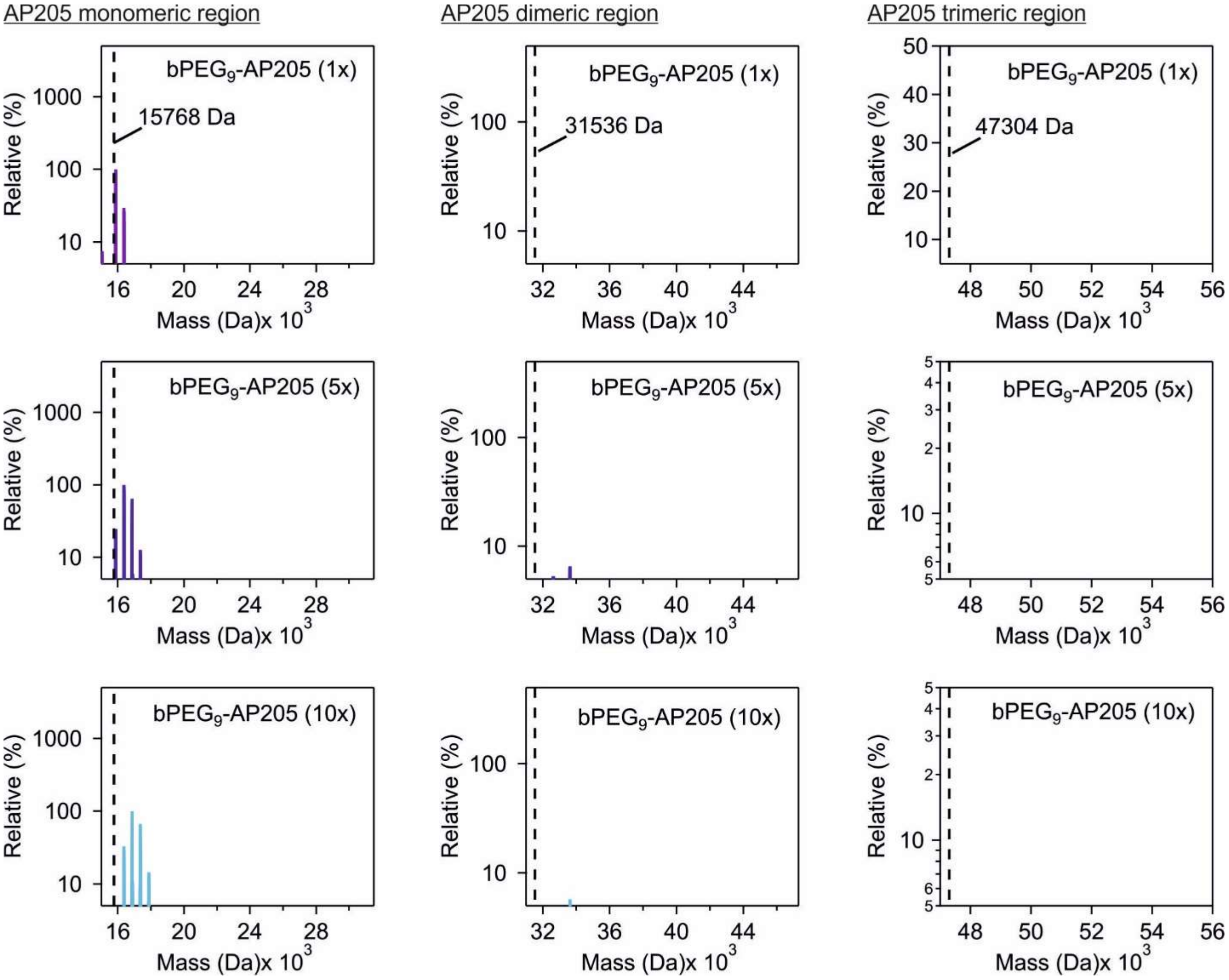

Figure S4-2: ESI-MS data of bPEG$_9$-AP205 VLP.

Table S4-1: Summary of the number and relative abundance of PEG$_9$-conjugated AP205 VLPs.

| Virus-like particle | Molar ratio[1] | 15768 Da + 0x MW of crosslinker | 15768 Da + 1x MW of crosslinker | 15768 Da + 2x MW of crosslinker | 15768 Da + 3x MW of crosslinker | 15768 Da + 4x MW of crosslinker | 15768 Da + 5x MW of crosslinker | 15768 Da + 6x MW of crosslinker | 15768 Da + 7x MW of crosslinker | 15768 Da + 8x MW of crosslinker |
|---|---|---|---|---|---|---|---|---|---|---|
| bPEG$_9$-AP205 | 1x | 15768 Da (100%) | 16265 Da (29.5%) | | | | | | | |
| bPEG$_9$-AP205 | 5x | 15768 Da (24.5%) | 16265 Da (100%) | 16761 Da (64.3%) | 17258 Da (12.7%) | | | | | |
| bPEG$_9$-AP205 | 10x | | 16264 Da (33.1%) | 16761 Da (100%) | 17257 Da (67.4%) | 17772 Da (14.6%) | | | | |

[1] [molecule/AP205 monomer]

#### ESI-MS of bPEG$_{13}$-AP205 VLP

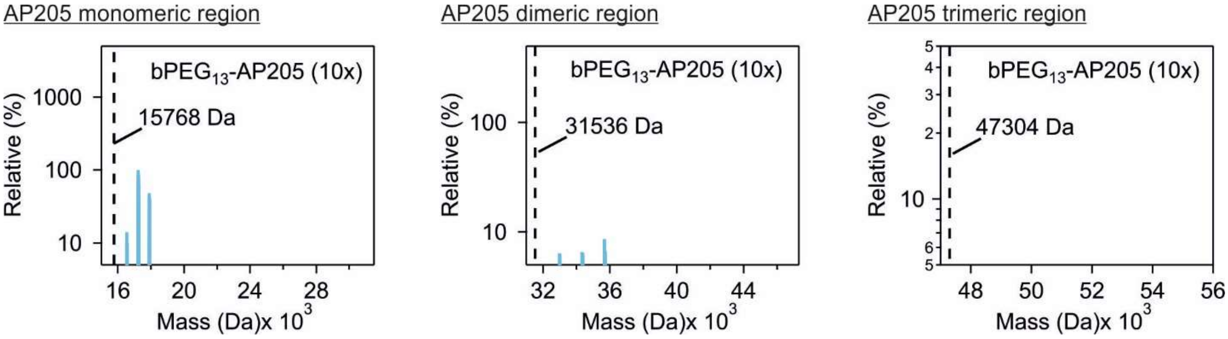

Figure S4-3: ESI-MS data of bPEG$_{13}$-AP205 VLP.

Table S4-2: Summary of the number and relative abundance of PEG$_{13}$-conjugated AP205 VLPs.

| Virus-like particle | Molar ratio[1] | 15768 Da + 0x MW of crosslinker | 15768 Da + 1x MW of crosslinker | 15768 Da + 2x MW of crosslinker | 15768 Da + 3x MW of crosslinker | 15768 Da + 4x MW of crosslinker | 15768 Da + 5x MW of crosslinker | 15768 Da + 6x MW of crosslinker | 15768 Da + 7x MW of crosslinker | 15768 Da + 8x MW of crosslinker |
|---|---|---|---|---|---|---|---|---|---|---|
| bPEG$_{13}$-AP205 | 10x | | 16441 Da (14.2%) | 17113 Da (100%) | 17786 Da (48.2%) | | | | | |

[1] [molecule/AP205 monomer]

#### ESI-MS of bPEG$_{17}$-AP205 VLP

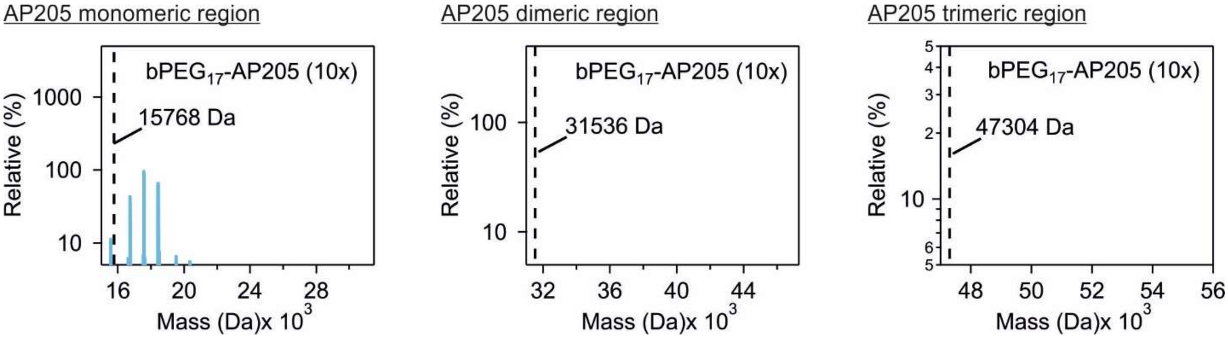

Figure S4-4: ESI-MS data of bPEG$_{17}$-AP205 VLP.

Table S4-3: Summary of the number and relative abundance of PEG$_{17}$-conjugated AP205 VLPs.

| Virus-like particle | Molar ratio[1] | 15768 Da + 0x MW of crosslinker | 15768 Da + 1x MW of crosslinker | 15768 Da + 2x MW of crosslinker | 15768 Da + 3x MW of crosslinker | 15768 Da + 4x MW of crosslinker | 15768 Da + 5x MW of crosslinker | 15768 Da + 6x MW of crosslinker | 15768 Da + 7x MW of crosslinker | 15768 Da + 8x MW of crosslinker |
|---|---|---|---|---|---|---|---|---|---|---|
| bPEG$_{17}$-AP205 | 10x | | 16617 Da (45.1%) | 17466 Da (100%) | 18332 Da (68.9%) | | | | | |

[1] [molecule/AP205 monomer]

#### ESI-MS of bPEG$_{21}$-AP205 VLP

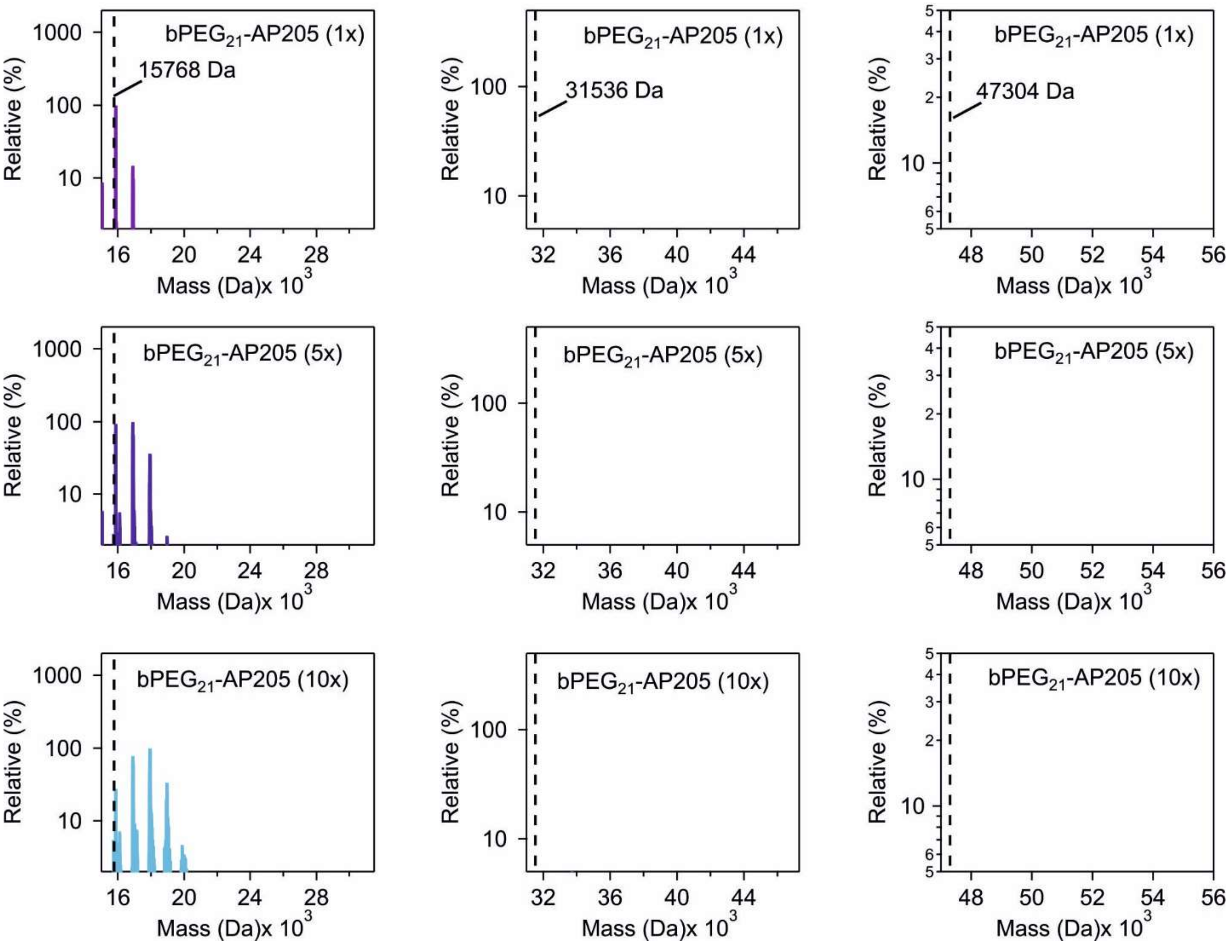

Figure S4-5: ESI-MS data of bPEG$_{21}$-AP205 VLP.

Table S4-4: Summary of the number and relative abundance of PEG$_{21}$-conjugated AP205 VLPs.

| Virus-like particle | Molar ratio[1] | 15768 Da + 0x MW of crosslinker | 15768 Da + 1x MW of crosslinker | 15768 Da + 2x MW of crosslinker | 15768 Da + 3x MW of crosslinker | 15768 Da + 4x MW of crosslinker | 15768 Da + 5x MW of crosslinker | 15768 Da + 6x MW of crosslinker | 15768 Da + 7x MW of crosslinker | 15768 Da + 8x MW of crosslinker |
|---|---|---|---|---|---|---|---|---|---|---|
| bPEG$_{21}$-AP205 | 1x | 15768 Da (100%) | 16793 Da (14.8%) | | | | | | | |
| bPEG$_{21}$-AP205 | 5x | 15768 Da (94.3%) | 16793 Da (100%) | 17835 Da (36.7%) | | | | | | |
| bPEG$_{21}$-AP205 | 10x | 15768 Da (27.8%) | 16793 Da (78%) | 17836 Da (100%) | 18861 Da (33.4%) | | | | | |

[1] [molecule/AP205 monomer]

#### ESI-MS of bPEG$_{25}$-AP205 VLP

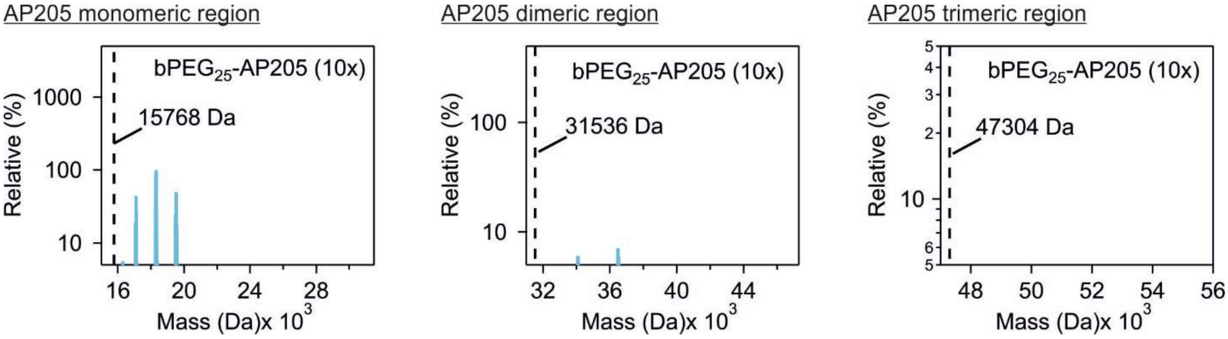

Figure S4-6: ESI-MS data of bPEG$_{25}$-AP205 VLP.

Table S4-5: Summary of the number and relative abundance of PEG$_{25}$-conjugated AP205 VLPs.

| Virus-like particle | Molar ratio[1] | 15768 Da + 0x MW of crosslinker | 15768 Da + 1x MW of crosslinker | 15768 Da + 2x MW of crosslinker | 15768 Da + 3x MW of crosslinker | 15768 Da + 4x MW of crosslinker | 15768 Da + 5x MW of crosslinker | 15768 Da + 6x MW of crosslinker | 15768 Da + 7x MW of crosslinker | 15768 Da + 8x MW of crosslinker |
|---|---|---|---|---|---|---|---|---|---|---|
| bPEG$_{25}$-AP205 | 10x | | 16987 Da (44.5%) | 18205 Da (100%) | 19406 Da (50.3%) | | | | | |

[1] [molecule/AP205 monomer]

#### Nano ultra-performance liquid chromatography coupled to mass spectroscopy (NanoUPLC-MS/MS) analysis of bPEG$_9$-AP205 VLP

The results showed that five out of seven lysine residues together with the N-terminus were modified by one-end conjugated BS(PEG$_9$) with the molecular formula $C_{22}O_{12}H_{40}$ (MW 496.3 Da). Furthermore, two “intra”-AP205 monomer linkages between the N-terminus (A2) and lysine 4 (K4) and between K94 and K101 were detected. In agreement with the results of reducing SDS PAGE, the LC-MS/MS analysis revealed the presence of “inter”-AP205 monomer crosslinking. In this case, two AP205 monomers were linked by doubly conjugated BS(PEG$_9$) with the molecular formula $C_{22}O_{11}H_{38}$ (MW 478.3 Da). The cross linkages were between K4 and K61 as well as between K4 and K94. More details in below:

It is shown in Table S4-6 that five out of seven lysine residues of AP205 monomer were modified by $C_{22}O_{12}H_{40}$, namely one-end conjugated BS(PEG$_9$). The raw data were analyzed by Byonic 4.0 (Protein Metrics, USA) with the consideration of mass tolerance of 5 ppm at MS1 level, and 20 ppm at MS2 level and a score higher than 200. To confirm the results, Spectrum Identification Machine (SIM-XL) was also used for crosslink analysis (*1*).

For crosslinked peptides, the results from SIM-XL showed that there were two intra-linked peptides at A2-K4 and K94-K101 (Figure S4-7 and Table S4-7). In addition, two cross-linked peptides were observed, namely A2/K4-K61 and A2/K4-K94. The software did not confirm either N-terminus A2 or lysine K4 participated in the crosslinking reaction with the other lysine residue (K61 or K94) due to the lack of key ions. However, if the major linkage was between N-terminus and K61 or K94, K4 residue would have been cleaved off by trypsin. Then, the results would have shown a linkage between $^{2}$AN$^{4}$K-crosslinked with the partner peptide. But the results showed $^{2}$AN$^{4}$KPMQPITSTAN$^{15}$K linked to the other peptide. Therefore, the data suggested that most likely, K4 instead of A2 was linked to the other lysine. Byonic results agreed with the assignment from the SIM-XL software.

The location of the afore-mentioned N-terminus and lysine residues are noted in the sequence below:

M.$^{2}$AN$^{4}$KPMQPITSTAN$^{15}$KIVWSDPTRLSTTFSASLLRQRV$^{38}$KVGIAELNNVSGQYVSVY$^{56}$KRPA
P$^{61}$KPEGCADACVIMPNENQSIRTVISGSAENLATL$^{94}$KAEWETH$^{101}$KRNVDTLFASGNAGLGFLD
PTAAIVSSDTTAGSGGAHATANATAHHHHHH

Table S4-6: Summary of Byonic analysis of bPEG$_9$-AP205 VLP based on the following criteria: score higher than 200 and peptides correctly cleaved at Lys and Arg residues.

| Position | Tryptic peptides | Observed linker[1] | Score | PSM |
|---|---|---|---|---|
| 2 or 4 | M.$^{2}$AN$^{4}$KPMQPITSTANK.I[2] | K4(+$C_{22}O_{12}H_{40}$) | 797.2 | 111 |
| 15 | M.AN$^{4}$KPMQPITSTAN$^{15}$KIVWSDPTR.L | K4(+$C_{22}O_{12}H_{40}$) K15 (+$C_{22}O_{12}H_{40}$) | 885.3 | 5 |
| 38 | R.V$^{38}$KVGIAELNNVSGQYVSVYK.R | K38(+$C_{22}O_{12}H_{40}$) | 954.7 | 14 |
| 56 | R.VKVGIAELNNVSGQYVSVY$^{56}$K.R | --[3] | - | - |
| 61 | R.PAP$^{61}$KPEGCADACVIMPNENQSIR.T | K61(+$C_{22}O_{12}H_{40}$) | 511.8 | 6 |
| 94 | R.TVISGSAENLATL$^{94}$K.A | K94(+$C_{22}O_{12}H_{40}$) | 534.7 | 2 |
| 101 | R.TVISGSAENLATLKAEWETH$^{101}$K.R | ---[3] | - | - |

Note:

1. The chemical format for one-end conjugated PEG$_9$ adduct on lysine is $C_{22}O_{12}H_{40}$ and the exact mass is 496.26. For PEG$_9$-crosslinked peptides, the mass is the sum of two peptides plus 478.26 ($C_{22}O_{11}H_{38}$).
2. The MS/MS spectra could not confirm which amino acid was modified by PEG$_9$ due to the lack of b1 and b2 ions.
3. The lysine was not labeled with PEG$_9$.

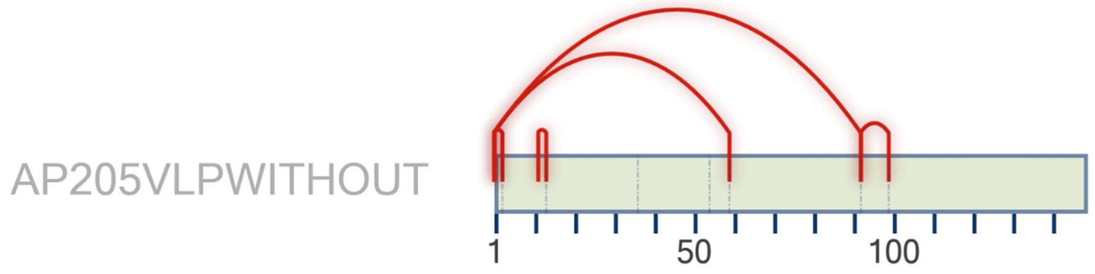

Figure S4-7: The scheme of crosslinked peptides from bPEG$_9$-AP205 VLP via SIM search engine. The fasta sequence that was used to perform the search started from A at the N-terminus. Therefore, the position of modified amino acids had 1 position less than the full sequence. The identified peptides are summarized in Table 2.

Table S4-7: Summary of $PEG_9$-cross-linked peptides from SIM-XL search engine

Intra-linked

| Scan Number | Experimental M+H | Type Peptide Link | Primary Score | ppm | Peaks Matched | Peptide Sequence | Position of XL-Residue 1 | Position of XL-Residue 2 |
|---|---|---|---|---|---|---|---|---|
| 14584 | 1979.02355 | | | | | | | |
| | | Intra | 2.97394 | 8.78 | 25 | ANKPMQPITSTANK | 1 | 3 |
| 14580 | 1979.022975 | | | | | | | |
| | | Intra | 2.81724 | 9.071 | 19 | ANKPMQPITSTANK | 1 | 3 |
| 14364 | 1979.02475 | | | | | | | |
| | | Intra | 2.72725 | 8.174 | 23 | ANKPMQPITSTANK | 1 | 3 |
| 19553 | 2919.51845 | | | | | | | |
| | | Intra | 2.25529 | 5.779 | 18 | TVISGSAENLATLKAEWETHKR | 93 | 100 |

Inter-linked

| Scan Number | Experimental M+H | Type Peptide Link | Primary Score | ppm | Peaks Matched | Peaks Matched ß | Peptide Sequence | Position of XL-Residue 1 | Position of XL-Residue 2 |
|---|---|---|---|---|---|---|---|---|---|
| 16148 | 4419.294875 | | | | | | | | |
| | | Inter | 5.59702 | 3.858 | 33 | 21 | TVISGSAENLATLKAEWETHKR - ANKPMQPITSTANK | 93 | 1 |
| | | Inter | 5.49283 | 3.858 | 33 | 20 | TVISGSAENLATLKAEWETHKR - ANKPMQPITSTANK | 93 | 3 |
| 12054 | 4688.3032 | | | | | | | | |
| | | Inter | 4.60746 | 3.46 | 31 | 12 | RPAPKPEGC(57.02146)ADAC(57.02146)VIMPNENQSIR - ANKPMQPITSTANK | 60 | 1 |
| | | Inter | 4.3861 | 3.46 | 31 | 10 | RPAPKPEGC(57.02146)ADAC(57.02146)VIMPNENQSIR - ANKPMQPITSTANK | 60 | 3 |
| 16128 | 4419.2897 | | | | | | | | |
| | | Inter | 3.85457 | 5.029 | 23 | 13 | TVISGSAENLATLKAEWETHKR - ANKPMQPITSTANK | 93 | 1 |
| | | Inter | 3.63072 | 5.029 | 23 | 11 | TVISGSAENLATLKAEWETHKR - ANKPMQPITSTANK | 93 | 3 |
| 12055 | 4688.3032 | | | | | | | | |
| | | Inter | 3.49869 | 3.46 | 24 | 9 | RPAPKPEGC(57.02146)ADAC(57.02146)VIMPNENQSIR - ANKPMQPITSTANK | 60 | 1 |

### S5. Characterization of PEGylated VLPs from the analysis of reducing SDS PAGE

The figure below shows the reducing SDS PAGE results of AP205 VLP, AP205 VLP in mock reaction buffer, $bPEG_9$-AP205 VLP, $bPEG_5$-AP205 VLP, BS3-AP205 VLP and Sulfo-NHS-AP205 VLP. The VLPs $bPEG_9$-AP205, $bPEG_5$-AP205, BS3-AP205 and Sulfo-NHS-AP205 were prepared at three molar ratios, 1x, 5x and 10x. In lane 1, the MW band in range 15–20 kDa corresponds to AP205 monomer. The same band appears in all the other lanes, expect that in lanes 5 and 6 versus 3 and lanes 7 and 8 versus 6, respectively for $bPEG_9$-AP205 VLP and $bPEG_5$-AP205 VLP, the bands are shifted to a higher molecular weight consistent with an increasing number of PEGylated AP205 monomers at higher molar ratio (c.f. S5). This statement does not apply to BS3-AP205 VLP and Sulfo-NHS-AP205 VLP which appear to be saturated already at 1x molar ratio. In cases of $bPEG_9$-AP205 VLP and $bPEG_5$-AP205 VLP, MW bands in range 25–37 kDa appeared and correspond to crosslinked AP205 dimers. Similarly, with $bPEG_9$-AP205 VLP and $bPEG_5$-AP205 VLP, MW bands around 50 kDa appeared and correspond to AP205 trimers. Bands corresponding to higher order oligomers are apparent for these VLPs. Unlike PEGylated VLPs, in lanes associated with BS3-AP205 VLP bands only in the dimeric region appeared while with Sulfo-NHS the band strength in the dimeric regions is similar to naked AP205 VLP.

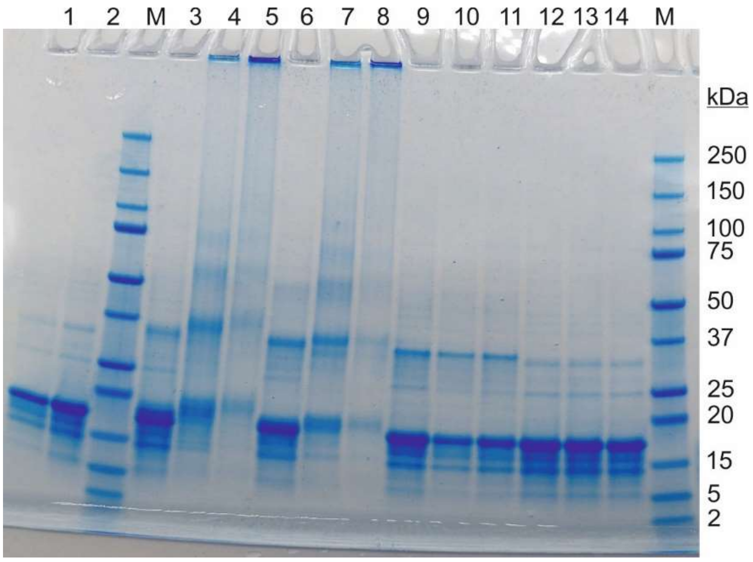

Figure S5-1: Reducing and denaturing SDS PAGE of AP205 VLP (lane 1), AP205 VLP in mock reaction buffer (lane 2), $bPEG_9$-AP205 VLP at three different molar ratios 1x, 5x and 10x (lanes 3–5 respectively), $bPEG_5$-AP205 VLP at molar ratios 1x, 5x and 10x (lanes 6–8 respectively), BS3-AP205 VLP at molar ratios 1x, 5x and 10x (lanes 9–11 respectively) and Sulfo-NHS-AP205 VLP at molar ratios 1x, 5x and 10x (lanes 12–14 respectively).

The figure below shows reducing SDS PAGE results of AP205 VLP, AP205 VLP in mock reaction buffer, $bPEG_5$-AP205 VLP, $bPEG_9$-AP205 VLP, $bPEG_{13}$-AP205 VLP, $bPEG_{17}$-AP205 VLP, $bPEG_{21}$-AP205 VLP, and $bPEG_{25}$-AP205 VLP. PEGylated VLPs were prepared at 1x molar ratio. The coat proteins of PEGylated VLPs showed a higher MW compared to AP205 coat protein monomer and the MW gradually increased with the length of the PEG. For $bPEG_{17}$-AP205 VLP (lane 6), $bPEG_{21}$-AP205 VLP (lane 7) and $bPEG_{25}$-AP205 VLP (lane 8), three to four bands were apparent in the monomeric MW range (15–25 kDa) that correspond to singly, doubly, triply, and quadruply PEG-conjugated AP205 monomers (c.f. S5). MW bands in the dimeric range (25–37 kDa) appeared for the PEG-crosslinked AP205 VLPs. Again, a clear graduation of PEG-crosslinked AP205 dimers bands corresponded to the MW of PEG crosslinkers. A dimer band is absent for AP205 VLP in lanes 1 and 2. PEG-crosslinked AP205 trimers and higher order oligomers are also apparent on the gel.

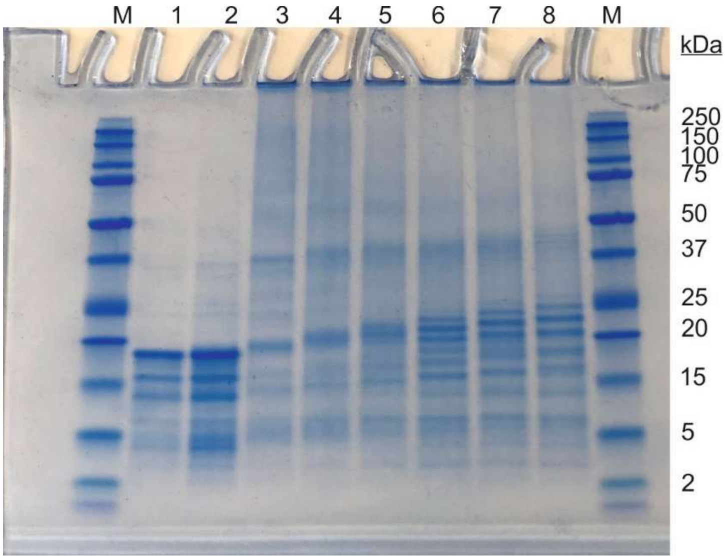

Figure S5-2: Reducing SDS PAGE of AP205 (lane 1), AP205 in mock reaction buffer (lane 2), $bPEG_5$-AP205 VLP (lane 3), $bPEG_9$-AP205 VLP (lane 4), $bPEG_{13}$-AP205 VLP (lane 5), $bPEG_{17}$-AP205 VLP (lane 6), $bPEG_{21}$-AP205 VLP (lane 7) and $bPEG_{25}$-AP205 VLP (lane 8). All preparations were at 10x molar ratio.

### S6. Investigation of colloidal stability

Using DLS, the colloidal stability of native and PEGylated AP205 VLPs were investigated in phosphate buffered saline (PBS) at pH 7.4, and in acidified PBS, pH 2.0–6.0. The pH-adjusted PBS was prepared by adding an appropriate amount of 100 mM HCl to PBS. Thereafter, 10 µl of 1 mg/ml VLP solution was added to 90 µl of acidified PBS. Since the addition of VLP solution increased the pH of acidified PBS, especially in the cases of pH 4.0 and pH 5.0, acidified PBS were prepared at a slightly lower pH. In summary, native AP205 VLP was stable at pH 7.4, but aggregated at pH values 2.0–6.0. In particular, already at pH 6.0, in the 'size distribution by intensity' spectrum, a peak associated with micron-size aggregates was observed. A pH dependent colloidal stability agrees with charge regulation of AP205 CPs. The PEGylation of AP205 VLP increased the stability profile up to pH ~ 4.0 mediated by steric repulsion of surface-conjugated PEG molecules and not by electrostatics. This is because the reaction of lysine residue or N-terminus with functional PEG linker reduces the positive charge of these moieties.

#### Native AP205 VLP

The colloidal stability of AP205 VLP was narrow, e.g., at pH ≤ 6.0, the VLP was readily aggregated. At pH 2.0, the VLP was somewhat stable. Therefore, the stable pH for AP205 VLP in PBS is pH 7.4.

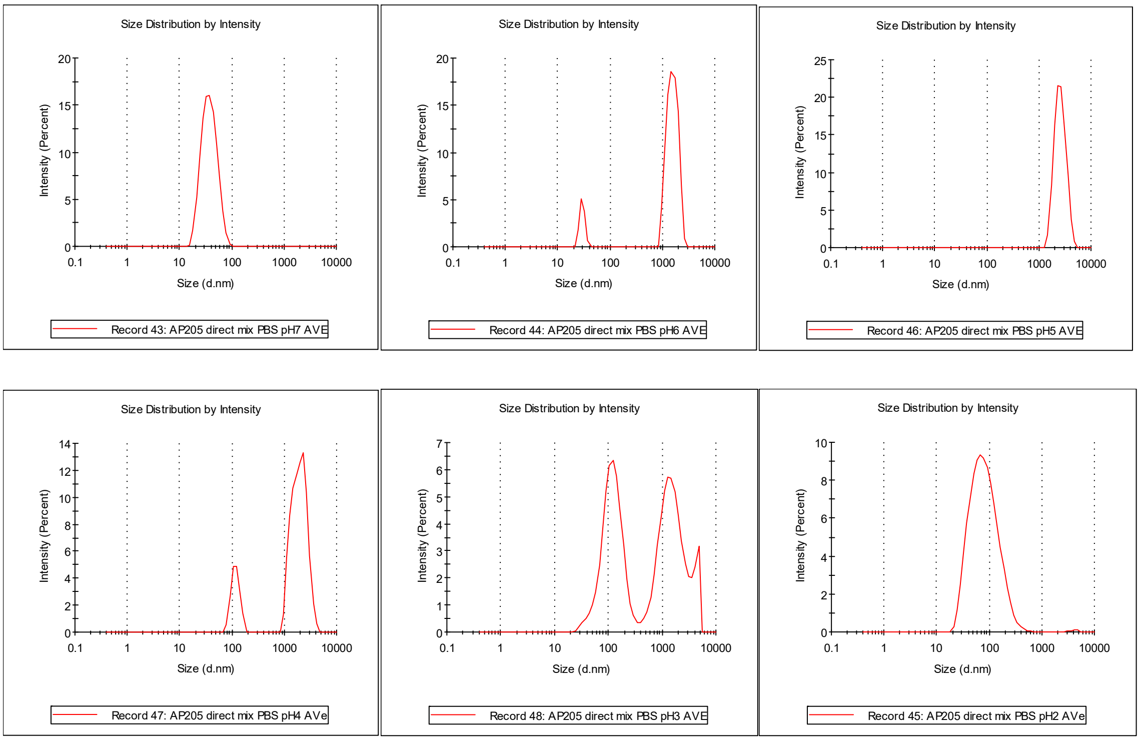

Figure S6-1: Hydrodynamic diameter of AP205 VLP at varying pH 7.4 to 2.0.

Table S6-1: Summary of the measurements from Figure S6-1. Cumulant and Contin analysis algorithms were used for data analysis. $D_h$ is hydrodynamic diameter, and PDI polydispersity index.

| pH | Cumulant $D_h$ (nm) | PDI | Contin (1st peak) $D_h$ (nm) |
|---|---|---|---|
| 7.4 | 35.0 | 0.12 | 38.8 |
| 6.0 | 1721 | 0.73 | 29.7 |
| 5.0 | 2470 | 0.21 | 2613 |
| 4.0 | 1458 | 0.88 | 118.3 |
| 3.0 | 268.4 | 0.89 | 127.5 |
| 2.0 | 68.5 | 0.23 | 93.5 |

**Comparison of colloidal stability between native and PEG-crosslinked AP205 VLPs**

The figure below shows the effect of pH on colloidal stability of native and PEG-crosslinked AP205 VLPs. The data suggested that AP205 VLP was aggregated in pH 6.0–3.0. The origin of this aggregation is electrostatics, i.e., as a function of pH, the surface charge density of AP205 coat protein monomers varies leading to instability of the VLP. In the cases of bPEG$_9$-AP205 VLP and bPEG$_{21}$-AP205 VLP, the VLPs remained stable up to pH 4.0. Therefore, PEG-crosslinking increased the colloidal stability at low pH.

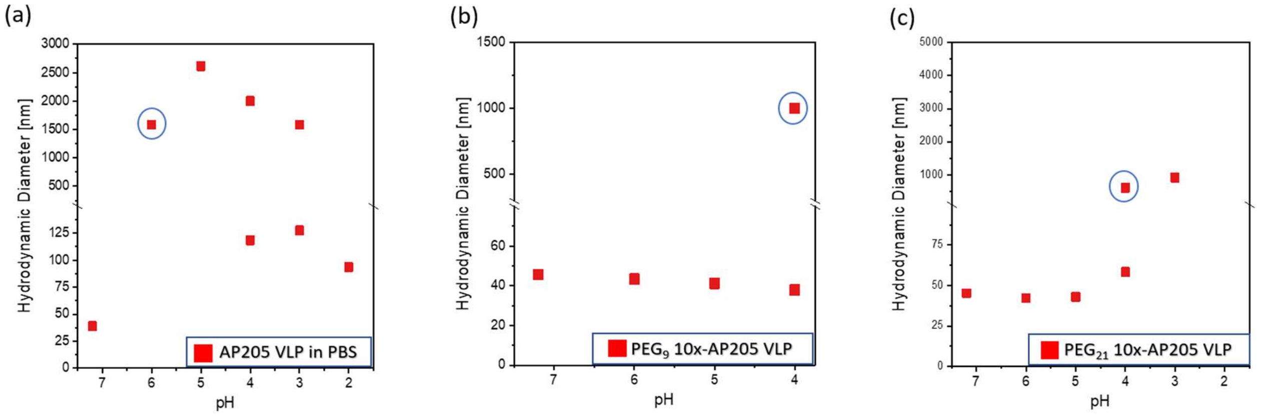

Figure S6-2: Hydrodynamic diameter of AP205 VLP, bPEG$_9$-AP205 VLP and bPEG$_{21}$-AP205 VLP at varying pH 7.4 to 2.0. Blue circles indicate the pH onset of aggregation from basic to acid solution.

### S7. Investigation of enzymatic stability in gastric fluid

Table S7-1: Type and pH of gastric fluid used in the investigations of enzymatic stability of naked and PEGylated AP205 VLPs.

| Type | Animal number | pH |
|---|---|---|
| Pig gastric fluid | 1314 | 3.0 |
| Pig gastric fluid | 1309 | 4.6 |
| Pig gastric fluid | 1257 | 5.5 |
| Mouse gastric fluid | Pooled | 4.6 |

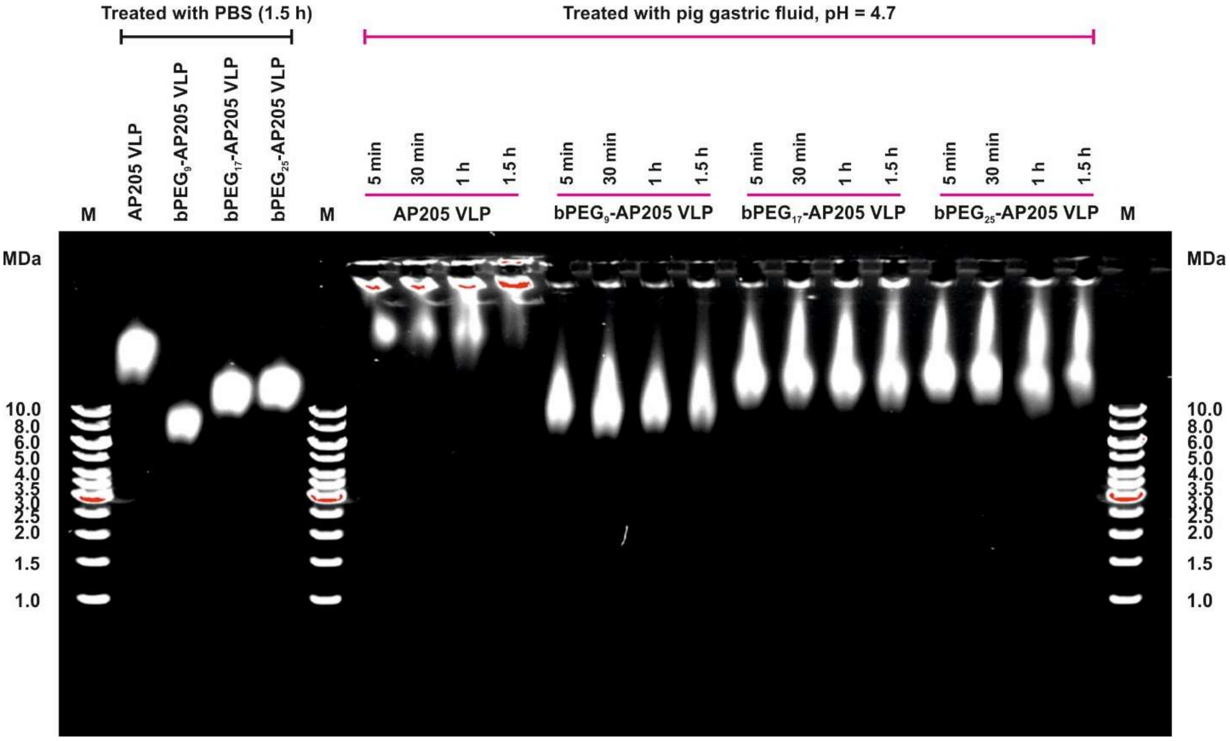

Figure S7-1. Enzymatic stability of naked and PEGylated VLPs in pig gastric fluid at pH 4.7. Agarose gel of AP205 VLP, $bPEG_9$-AP205 VLP, $bPEG_{17}$-AP205 VLP and $bPEG_{25}$-AP205 VLP in PBS and in pig gastric fluid, respectively on the left and right of the gel. VLP incubation times included 5 min, 30 min, 1 hour and 1.5 hours at 37°C. M stands for marker.

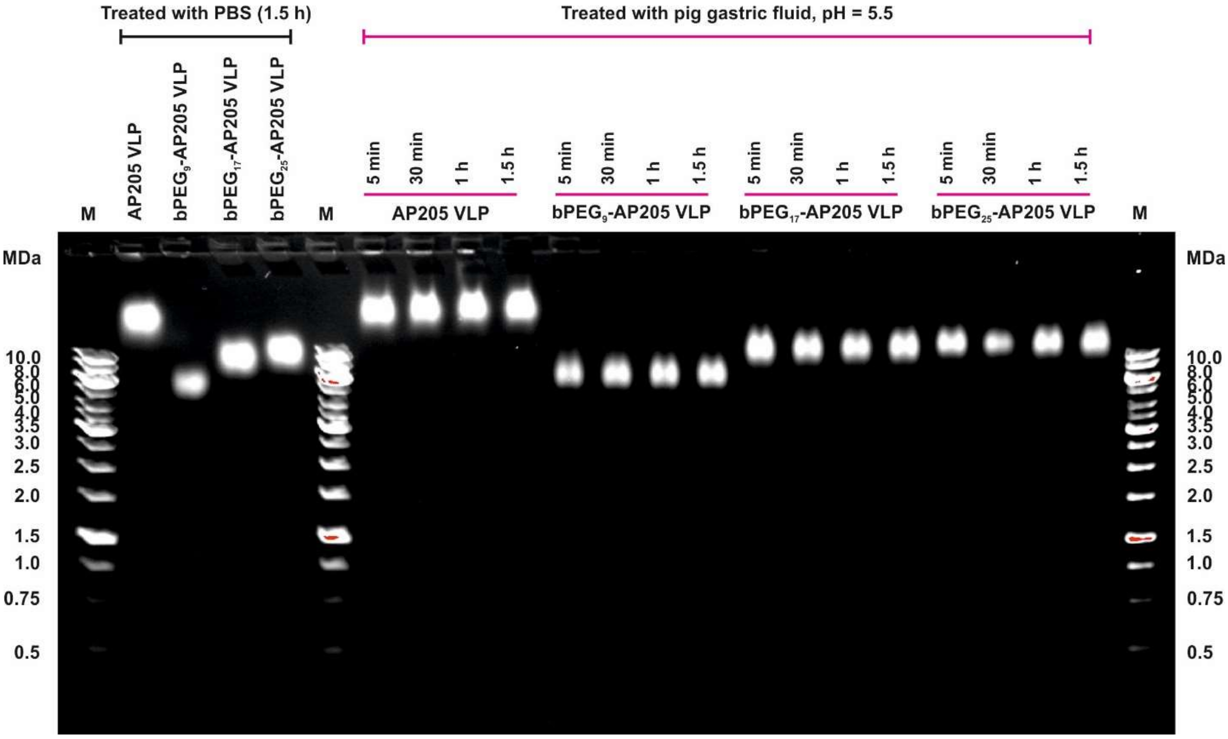

Figure S7-2. Enzymatic stability of naked and PEGylated VLPs in pig gastric fluid at pH 5.5. Agarose gel of AP205 VLP, bPEG$_9$-AP205 VLP, bPEG$_{17}$-AP205 VLP and bPEG$_{25}$-AP205 VLP in PBS and in pig gastric fluid, respectively on the left and right of the gel. VLP incubation times included 5 min, 30 min, 1 hour and 1.5 hours at 37°C. M stands for marker.

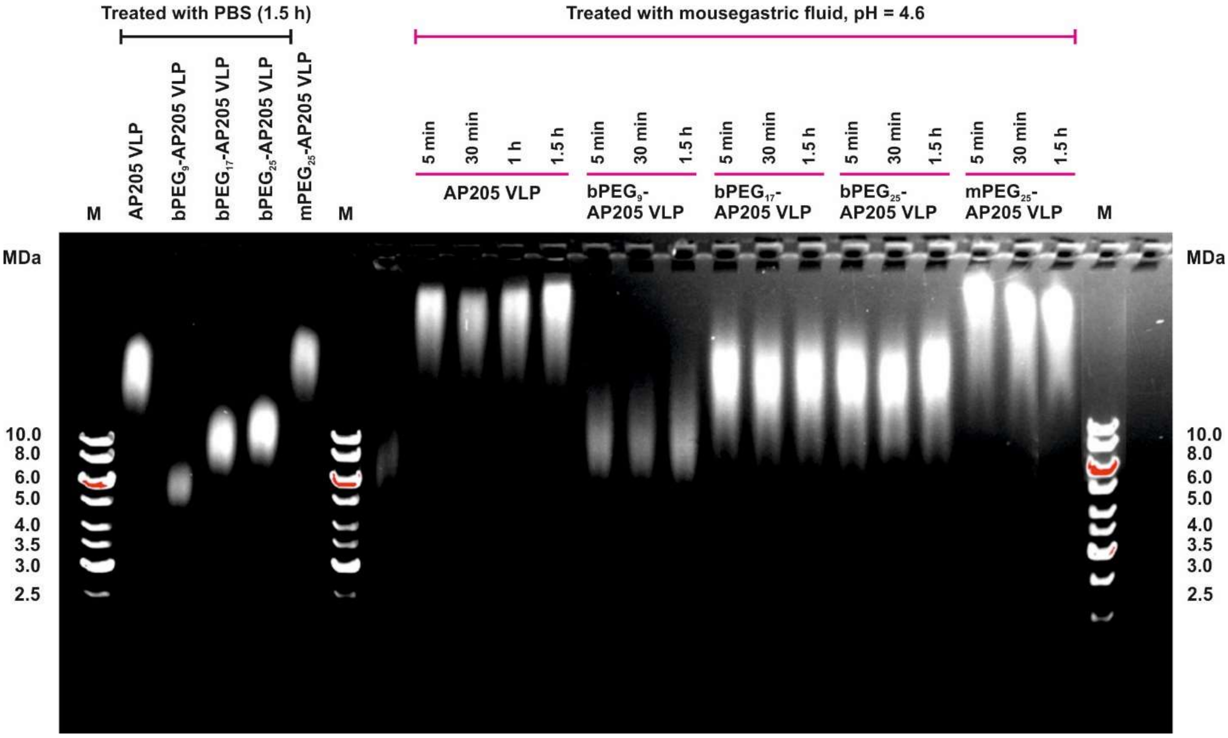

Figure S7-3. Enzymatic stability of naked and PEGylated VLPs in mouse gastric fluid at pH 4.6. Agarose gel of AP205 VLP, $bPEG_9$-AP205 VLP, $bPEG_{17}$-AP205 VLP, $bPEG_{25}$-AP205 VLP and $mPEG_{25}$-AP205 VLP in PBS and in mouse gastric fluid, respectively on the left and right of the gel. VLP incubation times included 5 min, 30 min and 1.5 hours at 37°C. M stands for marker.

### S8. Translocation of AP205 VLPs through mucus in *in vitro* 3D human nasal tissue

##### Results of VLP delivery to tissues 1 and 2

Figure S8-1 shows the translocation height and the mean fluorescence intensity (MFI) of the translocated native and PEG-crosslinked AP205 VLPs on the nasal monodonor tissue 1 (MD0860) and tissue 2 (MD0871). On these tissues, a similar accumulation Height is then found between the native AP205 VLP and the PEG-crosslinked AP205 VLPs was expected for the same tissue. However, the differences between the translocated MFIs are associated with the different permeability as well as adhesion to mucus and sedimentation. The general behavior shows that the VLPs behave somewhat similarly in terms of mucus translocation.

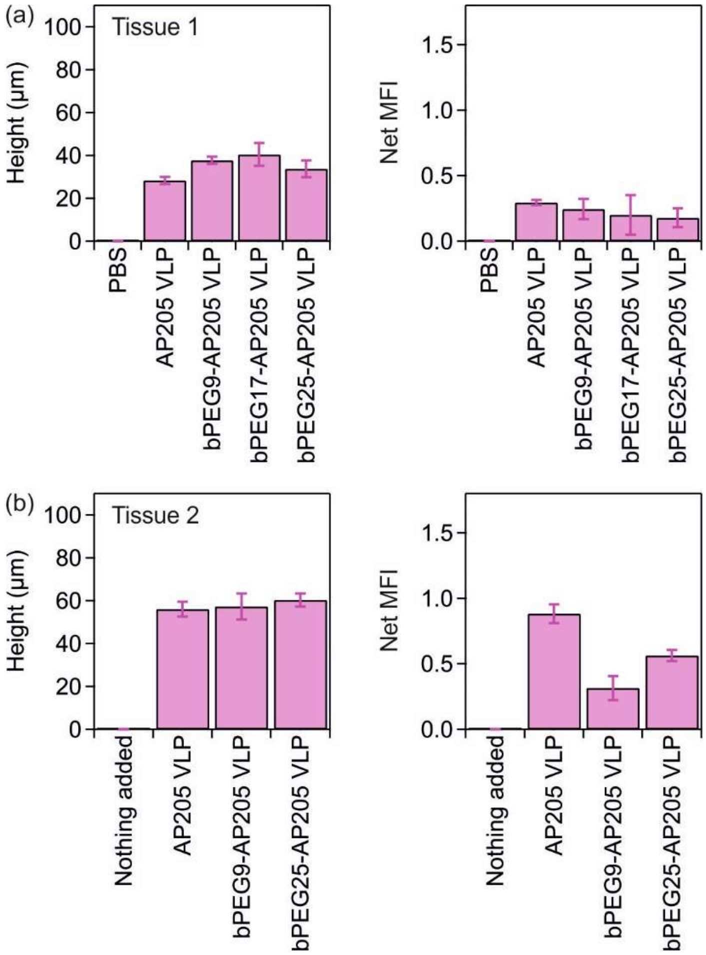

Figure S8-1: The translocation height and the mean fluorescence intensity (MFI) of the translocated native and PEG-crosslinked AP205 VLPs on nasal monodonor tissue 1 (MD0860) (a) and on nasal monodonor tissue 2 (MD0871) (b). In the experiments with tissue 1, PBS was added as a control. In the experiments with tissue 2, no liquid was added as control.

**Effect of MCC inhibition on lateral distribution and vertical translocation of VLPs**

Additional figures on the effect of ciliary beating in lateral distribution of delivered particles are shown below. In these experiments, tissue 3 (MD0774) was used.

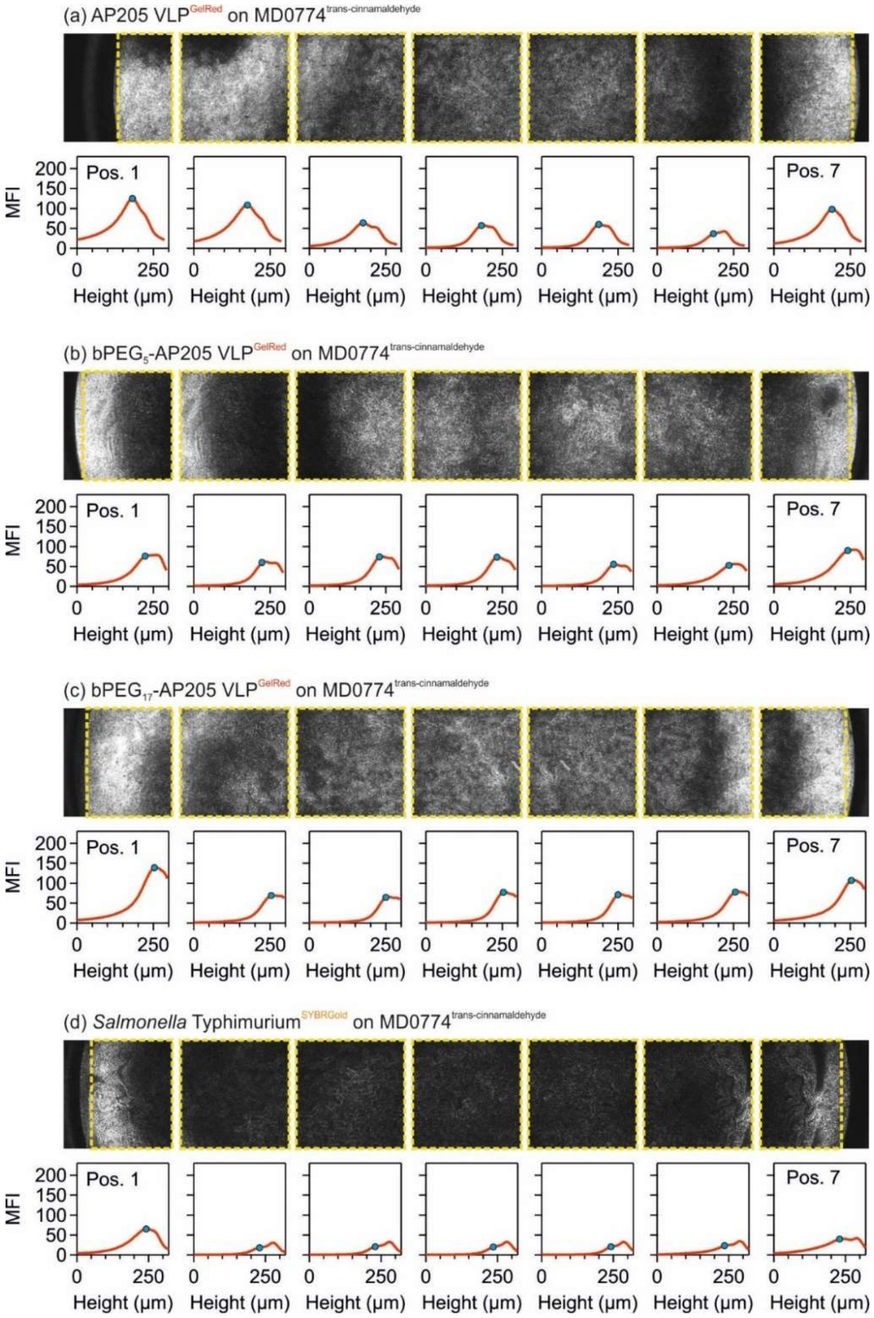

Figure S8-2. Effect of ciliary beating on distribution of delivered VLPs. Delivery of (a) AP205 VLP, (b) bPEG$_5$-AP205 VLP, (c) bPEG$_{17}$-AP205 VLP and (d) *Salmonella* Typhimurium to tissue 3 (MD0774) in the presence of ciliary beating inhibitory molecule trans-cinnamaldehyde. The yellow box denotes the area of mean fluorescence intensity (MFI) calculation and blue circle the location of VLP accumulation. In these figures, height starts from above the air-liquid interface (ALI) and increases in the direction of cell layer to below the membrane.
